# The Sequenced Genomes of Non-Flowering Land Plants Reveal the (R)Evolutionary History of Peptide Signaling

**DOI:** 10.1101/2020.06.02.130120

**Authors:** Chihiro Furumizu, Anders K. Krabberød, Marta Hammerstad, Renate M. Alling, Mari Wildhagen, Shinichiro Sawa, Reidunn B. Aalen

## Abstract

An understanding of land plant evolution is a prerequisite for in-depth knowledge of plant biology. Here we extract and explore information hidden in the increasing number of sequenced plant genomes, from bryophytes to angiosperms, to elucidate a specific biological question – how peptide signaling evolved. To conquer land and cope with changing environmental conditions, plants have gone through transformations that must have required a revolution in cell-to-cell communication. We discuss peptides mediating endogenous and exogenous changes by interaction with receptors activating intracellular molecular signaling. Signaling peptides were discovered in angiosperms and operate in tissues and organs like flowers, seeds, vasculature, and 3D meristems that are not universally conserved across land plants. Nevertheless, orthologues of angiosperm peptides and receptors have been identified in non-flowering plants. These discoveries provoke questions regarding the co-evolution of ligands and their receptors, and whether *de novo* interactions in peptide signaling pathways may have contributed to generate novel traits in land plants. The answers to such questions will have profound implications for the understanding of evolution of cell-to-cell communication and the wealth of diversified terrestrial plants. Under this perspective we have generated, analyzed and reviewed phylogenetic, genomic, structural, and functional data to elucidate the evolution of peptide signaling.

## INTRODUCTION

Land plants have evolved from freshwater green algae that started to colonize land ~ 470 million years ago (Mya) (Delwiche and Cooper, 2015; Ishizaki, 2017). This successful evolution has required groundbreaking biological innovations, such as the transition from the ancestral algal life cycle of haploid multicellular plants with zygotic meiosis to that with multicellular alternating haploid and diploid generations. While the diploid sporophyte and the haploid gametophyte initially might have employed the same genes, gene duplications followed by neofunctionalization likely facilitated the development of new specialized organs that necessitated extensive changes in gene expression patterns (Ligrone et al., 2012; Bowman et al., 2017; Jill Harrison, 2017) (Figure 1). Lycophytes and ferns, which appeared ~430 Mya (Kenrick and Crane, 1997; Gensel, 2008; Steemans et al., 2009; Morris et al., 2018), are extant plants representing early diverging lineages with vasculature, which provided mechanical support and transport of nutrients (Ishizaki, 2017). Roots evolved independently in lycophytes and euphyllophytes (vascular plants excepting lycophytes); they provide anchorage for the plants and allow acquisition of water and nutrients from the soil (Raven and Edwards, 2001; Hetherington and Dolan, 2019). The evolution of novel cell types, tissue types, and organs with increasing multicellular complexity must have put a high demand on cell-to-cell communication in land plants (Grienenberger and Fletcher, 2015).

**Figure 1.**
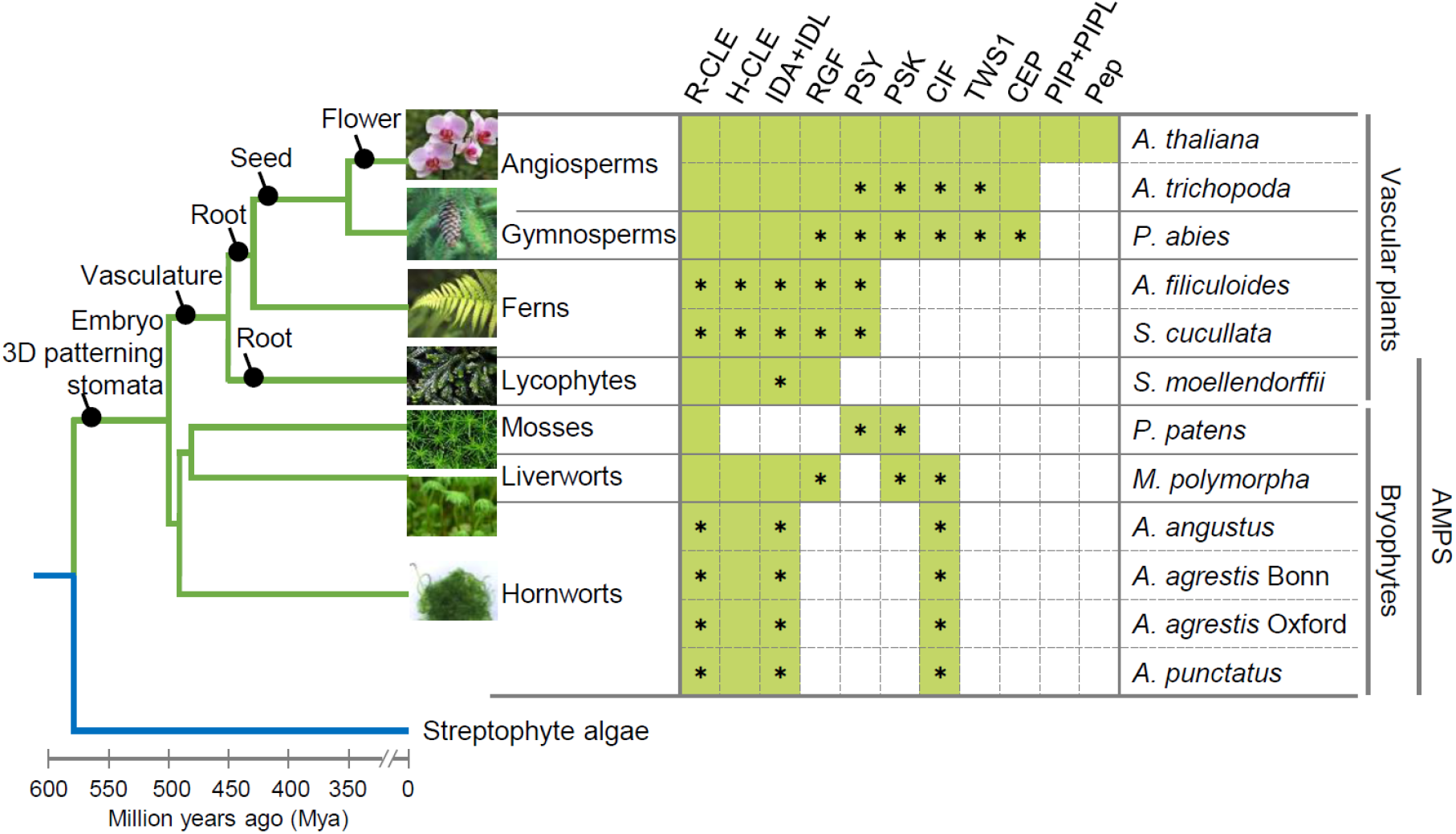
Phylogenetic distribution of small post-translationally modified peptides in land plants. **Left:** Simplified phylogenetic tree of major extant land plant lineages (green lines) based on (Transcriptomes, 2019). The branch lengths are roughly proportional to the estimated divergence dates (Morris et al., 2018). Gains of key morphological innovations are shown as black circles mapped on the tree while some gain events are still under discussion. **Right:** Green-colored boxes indicate the presence of homologues of the indicated small post-translationally modified signaling peptides for the model plants *Arabidopsis thaliana* (Lamesch et al., 2012), *Amborella trichopoda* (Amborella, 2013) (Amborella, 2013), *Picea abies* (Nystedt et al., 2013), *Azolla filiculoide*s and *Salvinia cucullata* (Li et al., 2018), *Selaginella moellendorffii* (Banks et al., 2011), *Physcomitrium patens* (Lang et al., 2018), *Marchantia polymorpha* (Bowman et al., 2017), and *Anthoceros* hornworts (Li et al., 2020; Zhang et al., 2020a). Asterisks indicate putative peptides newly identified in our sequence analyses. Images represent each taxa and are from Keiko Sakakibara (*A. agrestis* Oxford) and Pixabay (https://pixabay.com), except for *M. polymorpha*.

Recently, genomes of several hornwort *Anthoceros* species, the liverwort *Marchantia polymorpha,* and ferns (*Azolla filiculoides* and *Salvinia cucullata*) have been sequenced (Bowman et al., 2017; Li et al., 2018; Li et al., 2020; Yang et al., 2020; Zhang et al., 2020a). Together with the pioneering genome projects of the moss *Physcomitrium* (previously *Physcomitrella* (Rensing et al., 2020)) *patens*, the lycophyte *Selaginella moellendorffii*, the gymnosperm Norway spruce (*Picea abies*), and *Amborella trichopoda* representing the sister lineage to all other flowering plants, these resources have facilitated large-scale comparative genomic studies, which help us understand the evolution of land plants (Rensing et al., 2008; Banks et al., 2011; Amborella Genome, 2013; Nystedt et al., 2013). Advances in whole genome sequencing in streptophyte algae further provide opportunities to elucidate evolutionary processes of plant terresteralization (Nishiyama et al., 2018; Cheng et al., 2019; Chen et al., 2020; Jiao et al., 2020). For instance, the ancestry of several hormone and stress signaling pathways in land plants has been traced back to algae (Bowman et al., 2017; de Vries et al., 2018), underscoring the fundamental importance of these pathways for vegetative life.

Meanwhile, in flowering plants (angiosperms) the position of the classical hormones as the major mediators of signaling processes has over the last decades been challenged by small peptides interacting with leucine-rich repeat ectodomains of plasma-membrane-bound receptors with cytoplasmic serine/threonine kinase domains (KDs) (Matsubayashi, 2014; Olsson et al., 2019). Traditional hormones work over longer distances giving a so-called endocrine effect, while most peptides work over short distances and seem ideal for communication between neighboring cells with different functions (Wheeler and Irving, 2010). Changes in paracrine effects are likely to be instrumental in producing functional novelty or increasing complexity in an organism.

The first small signaling peptide was discovered in plants thirty years ago (Pearce et al., 1991), while more than 200 genes encoding leucine-rich repeat receptor-like kinases (LRR RLKs) were identified ten years later in the sequenced genome of *Arabidopsis thaliana* (Shiu and Bleecker, 2001b). They were assigned to fifteen subfamilies according to their unique structure, organization, and number of LRRs. Since then the increase in the number and quality of sequenced genomes, and advances in methods for *in silico* annotation of small genes, have together allowed the identification of hundreds of potential signaling peptides (Gong et al., 2002; Ghorbani et al., 2015; Tavormina et al., 2015; Goad et al., 2017). The peptides with known receptors represent two major groups, the cysteine-rich peptides (CRPs) (Supplemental Table 1), which attain their 3D structure by disulfide bridges formed between pairs of cysteines, and the post-translationally modified peptides (PTMPs, Table 1 and Supplemental Table 2), generated from prepropeptides and processed to mature peptides of 5-20 amino acids (Matsubayashi, 2011). To date, LRR-RLKs of the X, XI, and XIII subfamilies, with ligand-binding ectodomains built of >20 units of leucine-rich repeats, have been identified as main receptors for both groups of endogenous small signaling peptides. Here we will focus on the ancestry of the known PTMPs, which almost exclusively signal through the Arabidopsis receptor subfamily XI (hereafter LRXI).

**Table 1.**
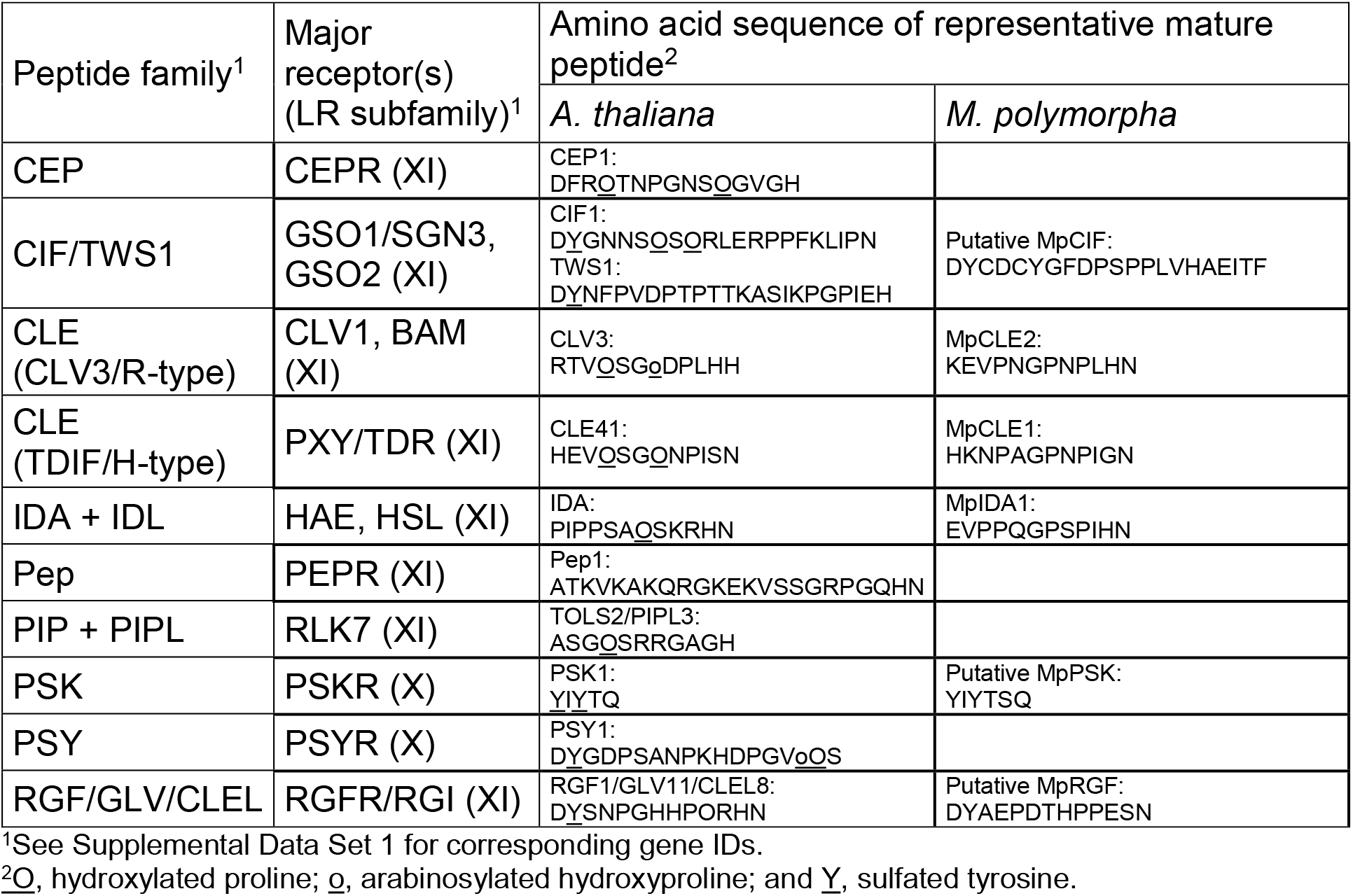
Major PTMP families in land plants.

LRR-RLKs have from their discovery been studied in a phylogenetic and evolutionary perspective (Shiu and Bleecker, 2001b; Lehti-Shiu et al., 2009; Liu et al., 2017; Chakraborty et al., 2019; Hosseini et al., 2020; Man et al., 2020). Signaling peptides have on the other hand been discovered due to their involvement in very “angiospermish” processes, like root growth, organ abscission, or seed development, and the phylogenetic investigations have focused on their distribution and function within the orders of flowering plants (Delay et al., 2013; Lori et al., 2015; Stø et al., 2015). Subsequent studies have matched LRR-RLKs with endogenous peptides as signaling pairs in angiosperms. Utilizing the increasingly available genome sequences of non-flowering plants, it is now feasible to elucidate peptide signaling in a phylogenetic and evolutionary perspective, taking into account the likely co-evolution of ligands and receptors.

LRXIs have been discovered in *M. polymorpha, P. patens* and *S. moellendorffii* (Bowman et al., 2017; Liu et al., 2017), but a detailed analyses of their relation to the well-studied Arabidopsis receptors has not been performed. Under the assumption that non-angiosperm LRXI receptors are activated by PTMP ligands, this discovery triggers questions regarding the presence and the roles of signaling peptides in different lineages of land plants. Changes in peptide signaling systems might be linked to cell-to-cell communication pathways that facilitated the evolution from ancestral algae to land plants and revolutionized terrestrial ecosystems. To address this hypothesis, we started by going through literature and data on peptide signaling and LRXI. A number of papers have reviewed specific peptide or receptor families involved in peptide signaling (Fernandez et al., 2013; Muschietti and Wengier, 2018; Oh et al., 2018; Taleski et al., 2018; Kaufmann and Sauter, 2019; Segonzac and Monaghan, 2019; Shi et al., 2019) and (Supplemental Table 2). It has not been, however, comprehensively investigated how widely these receptors and ligands are conserved; here we identified the need to search for homologues of Arabidopsis *LRXI* genes in non-flowering plants as well as genes encoding specific peptides to match the presence of their presumptive receptors. Thus, our *Perspective* reviews published data but is also supplemented with novel findings. All genes and proteins used are listed in Supplemental Dataset 1. We offer cross-comparison of multiple peptide ligand-receptor pairs and argue that if we are to understand the evolutionary history and possibly revolutionary importance of signaling peptides, we need to study cell-to-cell communication through peptide signaling pathways in an integrative manner, using diverse *in silico* methods that can generate hypotheses to be addressed by experimental approaches.

### PTMPs in land plants

PTMPs are generated from prepropeptides by post-translational processing and amino acid modifications (Matsubayashi and Sakagami, 2006; Olsson et al., 2019). Gene and genome duplications have led to evolutionary changes, but the amino acid sequences of C-terminal bioactive peptides have been conserved. Members of ten families of peptides with characteristic length and amino acid composition have been matched with LRR-RLKs (Table 1).

The CLAVATA3/EMBRYO SURROUNDING REGION-related (CLE) peptides have the broadest distribution, from bryophytes to angiosperms, and has largest family in Arabidopsis with 32 members (Jun et al., 2008; Goad et al., 2017). The 12-13 amino acids long CLE peptides are proline (Pro, P)-rich and characteristically have histidine-asparagine (His, H and Asn, N) or two His residues at the C-terminus (Table 1). CLE peptides are classified into two groups by the N-terminal residue: the R (arginine, Arg) and H groups. Many R-CLEs interact with CLAVATA1 (CLV1) or BARELY ANY MERISTEM (BAM) receptors involved in meristem maintenance in Arabidopsis (DeYoung et al., 2006; Hazak and Hardtke, 2016) while the three H-CLE peptides interact with the PHLOEM INTERCALATED WITH XYLEM/TDIF RECEPTOR (PXY/TDR) involved in vascular differentiation in Arabidopsis (Hirakawa et al., 2010). The CLE peptides are the first PTMPs from bryophytes that have been assigned with biological functions. In *M. polymorpha* an R-type peptide (MpCLE2) functions in a paracrine manner as a haploid stem cell-promoting signal in dichotomous branching of the thallus and is genetically dependent on the MpCLV receptor (Hirakawa et al., 2020).

The H-type peptide MpCLE1, with an MpTDR receptor, is on the other hand confined to the proliferative activity of gametophytic meristem and affects the overall size of reproductive organs (gametangiophore) (Hirakawa et al., 2019). In *P. patens* the R-type PpCLE peptide with its PpCLV receptor is involved 3D growth by determination of the orientation of stem cell division planes (Whitewoods et al., 2018). The function of the H-type CLE peptide recently detected in *Anthoceros* remains to be revealed (Li et al., 2020; Zhang et al., 2020b).

INFLORESCENCE DEFICIENT IN ABSCISSION (IDA) and the IDA-LIKE peptides are, similar to CLEs, 12-14 amino acid long P-rich peptides with C-terminal HN residues (Butenko et al., 2003; Jun et al., 2008; Vie et al., 2015) (Table 1). In Arabidopsis IDA signals trough HAESA (HAE) and HAE-LIKE2 (HSL2) to trigger cell wall remodeling leading to cell separation, in processes like floral organ abscission, lateral root emergence, and root cap sloughing (Cho et al., 2008; Stenvik et al., 2008; Butenko et al., 2014; Shi et al., 2018). Orthologues have been identified in all orders of flowering plants (Shi et al., 2019). Most recently, genes encoding putative IDA peptides and HSL receptors were identified in *M. polymorpha* (Bowman et al., 2017).

ROOT GROWTH FACTOR (RGF), also called GOLVEN (GLV) or CLE-LIKE (CLEL) peptides, are 12 amino acids long with Asp (Asp, D) and a sulfated tyrosine (Tyr, Y) at the N-terminus (Matsuzaki et al., 2010; Meng et al., 2012; Fernandez et al., 2013) (Table 1). Members of this family serve diverse roles in root development, like meristem homeostasis, gravitropism, and lateral root development (Fernandez et al., 2013). In addition to angiosperms, RGFs have been suggested to be present in the lycophyte *S. moellendorffii* (Ghorbani et al., 2015). RGFs can signal through RGF RECEPTOR (RGFR)/ROOT GROWTH FACTOR INSENSITIVE (RGI) receptors, which were discovered a few years ago (Shinohara et al., 2016; Song et al., 2016; Qiu et al., 2020).

When we initiated our study, the remaining peptide families of CASPAIAN STRIP INTERGRATING FACTORs (CIFs), CARBOXY-TERMINALLY ENCODED PEPTIDEs (CEPs), PAMP-INDUCED SECRETED PEPTIDEs (PIP/PIP-LIKE), PLANT ELICITOR PEPTIDES (Peps), Phytosulfokine (PSK), and PLANT PEPTIDEs CONTAINING SULFATED TYROSINE (PSY), Table 1) had not (yet) been identified in non-seed plants. However, the presence of putative *LRXI* (Bowman et al., 2017; Liu et al., 2017), raised the question of when signaling peptides evolved. This has not been well addressed in the literature; therefore we started to look for these peptide ligand homologues.

#### Methods for identification of peptide orthologues

A useful initial procedure for identification of homologues of Arabidopsis peptide-encoding genes is to search among more closely-related species, like monocots or *A. trichopoda*, using conventional Basic Local Alignment Search Tool for proteins (BLASTp) (Altschul et al., 1990) (https://blast.ncbi.nlm.nih.gov/Blast.cgi), confirm the homology of the hits with reciprocal BLASTp, and thereafter use identified peptides as queries in subsequent searches in gymnosperms and ferns, and later lycophytes and bryophytes. It is however, generally difficult to find distantly related peptide by BLASTp since prepropeptide sequences are highly variable except for the short peptide-encoding region. Small open reading frames encoding peptide precursors are additionally often overlooked during the annotation of newly sequenced genomes. Consequential false-negative problems may possibly be circumvented by using exhaustive tBLASTn searches against whole genome sequences or RNA collections translated in all six frames (Altschul et al., 1997).

BLASTp searches work more efficiently when limiting the query sequence to the region encompassing the mature peptide and using the BLAST-short option with the PAM30 matrix (Altschul et al., 1997). An alternative approach is to specify conserved motifs of a given peptide family in Pattern Hit Initiated (PHI) BLAST searches, and/or use queries with wildcards in positions where the amino acid residues are less conserved, e.g. we identify RGF homologues using the query DYxxxxxKPPIHN. PHI-BLAST with the DY motif facilitated also successful identification of *P. patens* PSK and PSKY1, and *M. polymorpha* CIF and PSK against the NCBI database (Zhang et al., 1998). The amino acids HN or HH, found at the C-terminus in several peptide families (Table 1), may also be used in PHI-BLAST. The likelihood of getting false-positive hits in BLAST searches increases with the size of the database searched, and it is therefore helpful to limit searches to coding sequences of less than 300 amino acids. This measure was employed to retrieve putative CIF and IDA peptides in *Anthoceros* species by the EMBOSS 6.5.7 tool *fuzzpro* (http://emboss.open-bio.org/rel/rel6/apps/fuzzpro.html). Queries with wildcards were generated based on consensus sequences from MUSCHLE alignments (Edgar, 2004) of mature peptides of Arabidopsis, *A. trichopoda, P. glauba, M. polymorpha* and *S. moellendorffii,* (DYxxxxPxPPLxxPxPF and PIPxSxPSKRHN, respectively) and used against protein libraries generated from gene-annotated *Anthoceros* genomes. Further options may be to test Machine Learning or Deep Learning methods that recently have been developed for peptide discovery (Plisson et al., 2020; Serrano, 2020; Zhang et al., 2020b).

The confidence of retrieved hits is usually low, and hits must therefore in all cases be analyzed for an appropriate prepropeptide length (<200 amino acids), the position of the assumed mature peptide sequence near or at the C-terminal end and the presence of a hydrophobic N-terminal secretion signal. When interpreting the sequences, it has to be kept in mind that sequences from extant species may have diversified from ancestral sequences.

Using these strategies above, we have identified putative peptide gene homologs in hornworts, lycophytes, ferns, and gymnosperms (Figure 1, indicated by asterisks, see Supplemental dataset 1), suggesting that orthologues of the respective interacting LRXI should be present.

### LRR-RLK subfamily XI in land plants

Shiu and Bleecker pioneered the characterization of RLKs in Arabidopsis and also initiated a comparison to other species (Shiu and Bleecker, 2001b; a). The total number of *LRR-RLK* genes in land plant species varies approximately from 60 to 400 in the genomes analyzed to date (Liu et al., 2017). The ectodomain is characteristic for each of the fifteen LRRs subfamilies, but the KD, as well as the scaffold amino acids of the LRRs ectodomain, are highly conserved. Therefore BLASTp searches of LRR-RLKs return many high-scoring hits. The challenge is to identify the few top hits that are true orthologues of the query protein: This can be achieved by reciprocal BLASTp searches in combination with phylogenetic analyses where paralogues will branch off outside the clade with the original query sequence. Phylogenetic analyses of all RLK subfamilies have therefore been based on the KD, often with limited taxon sampling (Zan et al., 2013; Wei et al., 2015; Liu et al., 2016; Magalhães et al., 2016; Zhou et al., 2016; Bowman et al., 2017; Lin et al., 2018; Man et al., 2020). This gives ample information on the relation between different subfamilies.

Our focus, however, is different; it is to explore the evolutionary history of the group LRXI receptors known to interact with signaling peptides in angiosperms. One of the most plausible hypotheses of the land plant phylogeny proposes an early separation of both monophyletic bryophyte and vascular plant lineages (Figure 1) (2019; Li et al., 2020; Zhang et al., 2020b). We have therefore sampled putative LRXI receptors from *<u>A</u>nthoceros* hornworts, the liverwort *M. polymorpha*, the moss *<u>P.</u> patens*, and the lycophyte *<u>S.</u> moellendorffii* (hereafter collectively referred to as AMPS), which are extant species representing lineages that arose from ancient splits in land plant diversification before the emergence of angiosperms.

#### Methods for phylogenetic analyses

In order to explore the origins and evolution of the group LRXI peptide-interacting receptors, we used 27 full-length receptors encoded by *LRXI* genes of Arabidopsis as individual queries in BLASTp searches against protein sequences from 11 species in the Phytozome database database (v13), the NCBI Genome database (https://www.ncbi.nlm.nih.gov/genome/), and the Hornworts database (https://www.hornworts.uzh.ch/en.html). These species were selected to cover the major evolutionary clades of land plants. In addition to the AMPS including a recently sequenced hornwort (*Anthoceros agrestis*) (Frangedakis et al., 2020; Li et al., 2020; Zhang et al., 2020a), we sampled *LRXIs* from banana, rice and maize monocots (*Musa acuminata, Oryza sativa,* and *Zea mays*), and three dicots, *Capsella rubella, Populus trichocarpa* and *Solanum lysopersicum* (pink Shepherd’s-purse, poplar, and tomato) (Supplemental data set 1). *A. trichopoda* was included in the analyses as a landmark for identification of evolutionary changes that took place before and after the emergence of angiosperms. Finally, we identified LRR-RLKs with more than 20 LRRs from the recently sequenced algae *Penium margaritaceum* (the Penium Genome Database (http://bioinfo.bti.cornell.edu/cgi-bin/Penium/home.cgi) (Jiao et al., 2020), *Spirogloea muscicola,* and *Mesotaenium endlicherianum* (Cheng et al., 2019) to be used as outgroup.

For each of the proteins encoded by the 27 Arabidopsis *LRXI* genes, we retrieved the 20 top BLASTp hits from each of the genomes. Duplicates and isoforms were removed before the candidate LRR sequences (1521 in total) were aligned with the l-INS-I algorithm in MAFFT v7.453 (Katoh and Standley, 2013). Ambiguously aligned characters were masked with trimAl (Capella-Gutiérrez et al., 2009) using the *gappyout* algorithm, and a preliminary phylogenetic tree was constructed with FastTree2 (Bennett and Scheres, 2010). This tree was used to identify sequences that did not belong to XI-type LRR-RLKs. The final dataset consisted of 249 sequences which were unmasked, and after annotation of domains with Interproscan5 (Jones, 2014) were split on each side of the transmembrane region to generate an LRR and a KD dataset. (Supplemental Dataset 1) Since the ectodomains of the LRXI have at least 20 LRRs and can be aligned, we took the opportunity to compare the ligand-interacting domain of these receptors, not only the KD. After separate realignment with MAFFT l-INS-I, and trimming with trimAl (*gappyout*, the LRR-containing ectodomain alignment was 954 amino acids long, and the KD alignment was 439 amino acids long (see Supplemental data sets 1, 2 and 3 for sequences and alingments).

Phylogenetic trees were constructed separately for each domain with IqTree (Supplemental Data Sets 2 and 3), and a tanglegram (Scornavacca et al., 2011) was drawn that connects the corresponding taxa in the two trees to help explore the differences in phylogenetic topology between the two domains of the LRXI proteins (Figure 2, Supplemental Figure 1). The tree files are available as Supplemental Data Sets 4 and 5.

**Figure 2.**
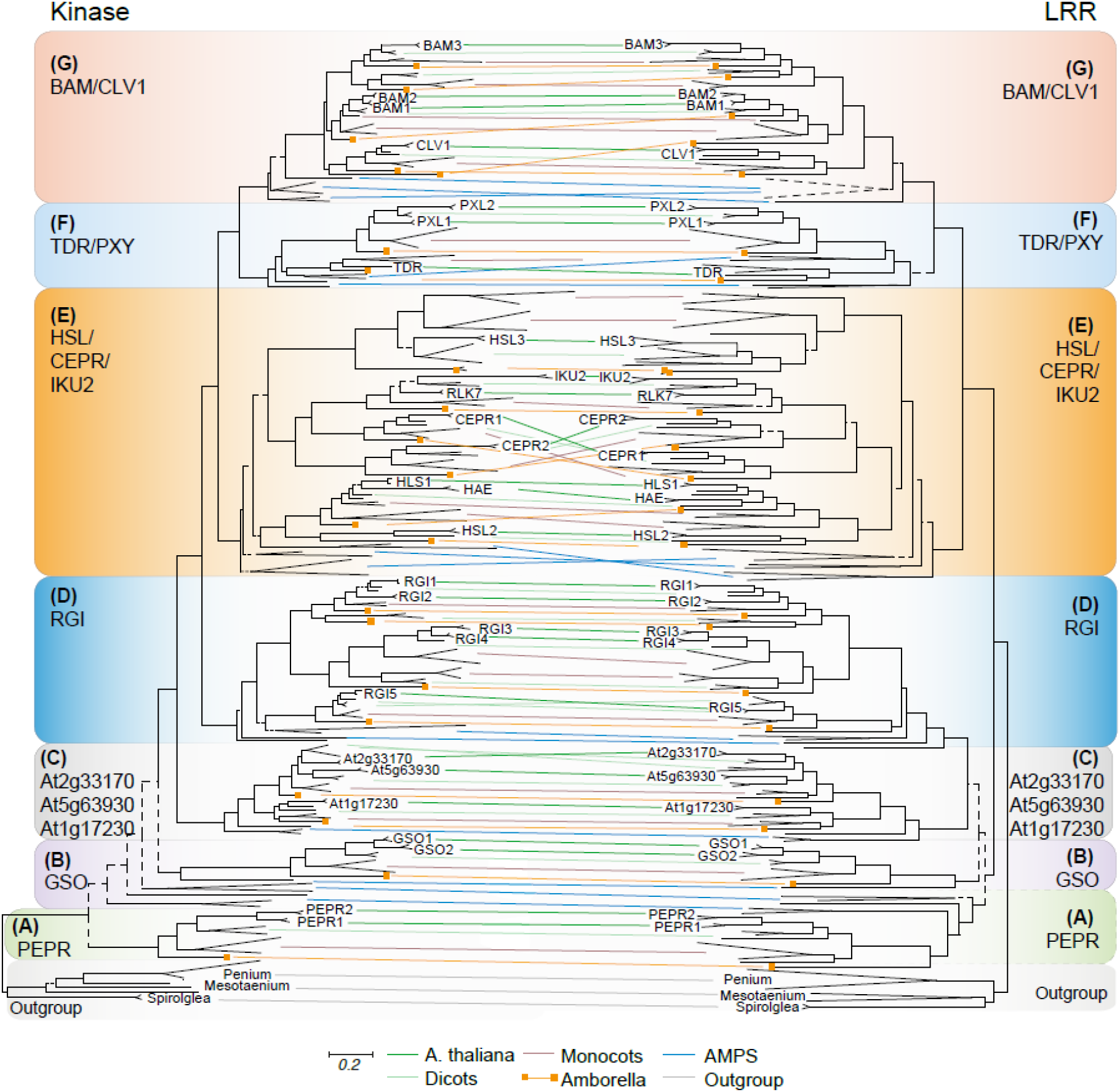
Tanglegram of the kinase domains (KDs) and the LRR ectodomains encoded by *LRXI* genes. **(A)-(G)** The seven colored boxes represent monophyletic clades of the indicated angiosperm LRXIs with a common origin and a last common ancestor with one or more AMPS (see Table 2 and Supplemental Figure 1 for details). The phylogenetic trees were constructed separately for the LRRs and the KDs. The braches representing Arabidopsis KDs or LRRs are labeled at the tips. A tanglegram connects KD and LRR originating from the same LRXI. The connecting lines represents *A. thaliana* (dark green); three dicot species (light green); three monocot species (brown); the basal angiosperm *A. trichopoda* (orange with boxes); the hornwort *Anthoceros,* the liverwort *M. polymorpha*, the moss *P. patens*, and the lycophyte *S. moellendorffii* (blue, collectively referred to as AMPS), and three species of Zygnematophyceae algae used as the outgroup (grey). Connecting lines between monophyletic groups of the same kind (i.e. either monocots, dicots, *A. trichopoda*, AMPS, or the outgroup) have been collapsed for clarity. For data on each species see Supplemental Figure 1. Branches drawn with solid lines are well supported (SH-aLRT > 0.85), while dashed lines represent unsupported branches with SH-aLRT < 0.85. Support values for all branches can be found in Supplemental Figure 1. The scale bar for branch lengths represents the average number of changes in amino acids per site, and the two trees of the tanglegram are drawn to scale.

In addition, nucleotide sequences of *LRXI* genes were aligned using Pal2Nal (Suyam et al., 2006), and the pairwise ratios of nonsynonymous to synonymous substitutions (Ka/Ks, also known as ω or dN/dS ratio) were calculated with the maximum likelihood method using PAML (Yang, 1997) for the two domains separately. These data were examined to investigate whether the selective pressure differs between the two domains of different functions: the ectodomain in perceiving peptide signals and the KD determining the subsequent intracellular response.

### Seven clades of LRXIs in land plants

The phylogenetic tanglegram consists of seven major monophyletic clades (Figure 2A-G, colored boxes), all with high support (Shimodaira-Hasegawa approximate likelihood ratio test (SH-alrt) > 85 (Shimodaira and Hasegawa, 1999); see Supplemental Figure 1 for all support values). However, the branching pattern differed between the KD phylogeny and the LRR phylogeny with regards to the putative orthologues of PEP RECEPTOR (PEPR), which in Arabidopsis play a role in innate immunity, and GASSHO (GSO), involved in the regulation of stem cell identity, control of seedling root growth, and establishment of the Casparian strip, which works as a diffusion barrier in the root vasculature (Tsuwamoto et al., 2008; Doblas et al., 2017; Nakayama et al., 2017 {Racolta, 2014 #5882) (Figure 2A-B). In the KD phylogeny, a paraphyletic group of AMPS other than *M. polymorpha* branches in between GSO and PEPR, while in the LRR phylogeny these AMPS form a monophyletic clade with PEPR. Except for one *M. polymorpha* orthologue of GSO, the phylogenetic trees could not unambiguously resolve the relationship between AMPS and angiosperm homologues as the support values for the deep nodes were low in both phylogenies, which reflects a common problem of inferring ancient relationships of *LRR-RLK* genes (Liu et al., 2016).

The remaining five LRXI clades (Figure 2C-G) are all monophyletic in both the KD and LRR phylogenies, with high support (SH-aLRT support higher or equal to 99) and include sequences from both vascular plants and bryophytes, indicating that these clades existed in the most recent common ancestor of vascular plants and bryophytes. One of these clades (Figure 2C) encompass receptors encoded by three Arabidopsis genes (*At2g33170*, *At5g63930*, and *At1g17230*) of common ancestry with *S. moellendorffii* and *P. patens* orthologues, which all three are without known ligand(s) and function(s). Multivalent mutants may be needed to obtain lines with informative mutant phenotypes.

The neighboring RGI clade includes sequences from *M. polymorpha* and *S. moellendorffii*. Arabidopsis RGIs are in particular involved in developmental processes in roots in Arabidopsis (Fernandez et al., 2013; Shinohara et al., 2016). Roots have been suggested to have evolved independently in several lineages (Fujinami et al., 2020), and it would be interesting to explore whether RGFs and RGIs have laid a fundament for root development. Non-AMPS sequences form three subgroups each of which consists of orthologues of Arabidopsis RGI5, RGI3 and 4, and RGI1 and 2 (Figure 2D, Table 2). In contrast, the monophyletic group encompassing the angiosperm HSL-related receptors (Figure 2E, Table 2) have orthologues from all AMPS.

**Table 2.**
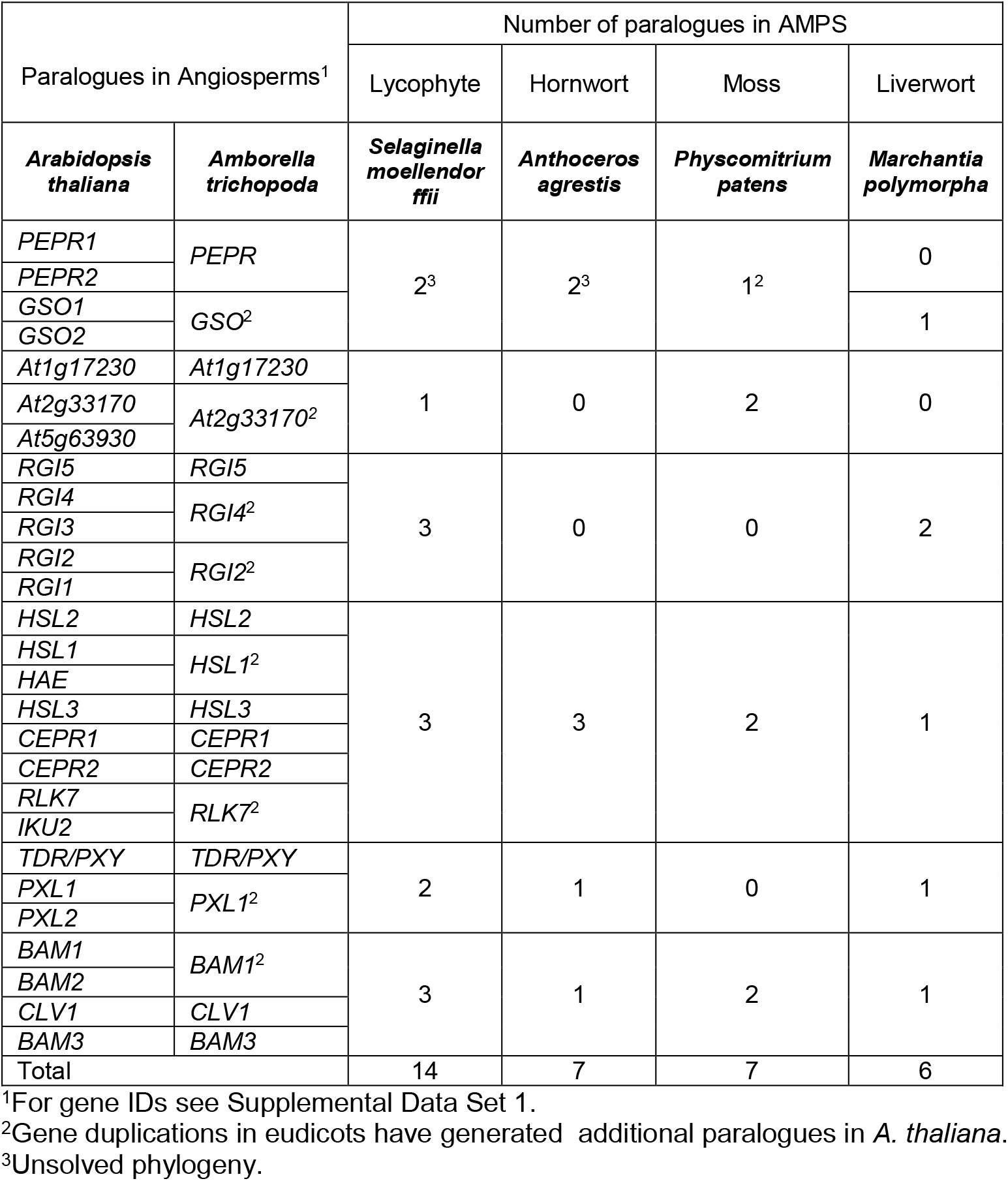
Presence of orthologues of Arabidopsis LRR-RLKs subfamily XI in *A. trichopoda* and AMPS.

The HSL clade has two major branches. One includes HAESA (HAE) and HSL2, which control cell separation events in Arabidopsis with the IDA peptide ligand (Shi et al., 2019). Consistent with a phylogenetic investigation of HSL receptors in angiosperms (Stø et al., 2015) HAE appeared during the evolution of dicot flowering plants. The other branch includes the Arabidopsis At5g25930 (also called HSL3) which very recently was shown to be involved in regulation of stomata closure by modulation of H2O2 levels in guard cells (Liu et al., 2020). This receptor and its orthologues are worth an investigation, as an ancestral *HSL3* gene has proliferated in several species, especially in monocots, both before and after the emergence of flowering plants (Figure 2E, Supplemental Figure 1). Branching off from HSL3, *A. trichopoda* orthologues of C-TERMINALLY ENCODED PEPTIDE (CEP) RECEPTORs (CEPRs), and RECEPTOR-LIKE KINASE7 (RLK7) are found (Figure 2E, Supplementary Figure 1). And finally, HAIKU2 (IKU2) branched off from RLK7 during eudicot evolution. CEPR mediates nitrogen starvation signaling (Tabata et al., 2014). RLK7 is involved in innate immunity and lateral root formation (Hou et al., 2014; Toyokura et al., 2019). *IKU2* of the most recent origin controls seed size in Arabidopsis (Luo et al., 2005) and may have been recruited to regulate angiosperm- or eudicot-specific characters (Friedman and Williams, 2004). So far the ligand of HSL3 and IKU2 is not known.

Interestingly, the two CEPR receptors have swapped places in the two trees; RLK7 and IKU2 branches off from CEPR1 in the KD tree but from CEPR2 in the LRR tree (Figure 2E, Supplemental Figure 1). Close inspection of the alignment of these four receptors indicates that CEPR1 has lost a repeat compared to the other three (Supplemental Figure 2). Interestingly, in the related HAE receptor, this repeat is involved in the interaction with the SERK1 co-receptor (Santiago et al., 2016). In the KD, RLK7 and IKU2 are more similar to CEPR1, and the CEPR2 is deviating, for instance with a truncated kinase motif M3 (Supplemental Figure 2). Thus, RLK7 and IKU2 may be more like CEPR2 with regards to perceiving signals, but more similar to CEPR1 with regards to output from the KD.

TDR/PXY and BAM/CLV receptors form clades distinct from the HSL. As mentioned above, these receptors interact with CLE ligands and are mainly involved in the regulation of stem cell activity. Consistent with previous findings, no TDR/PXY orthologues were found in *P. patens* (Figure 2F, Table 2), but other species tested have at least one sequence in the TDR/PXY clade. In its sister group, BAM/CLV1, we identified sequences from all AMPS (Figure 2G, Table 2) (Bowman et al., 2017; Liu et al., 2017; Li et al., 2020; Yang et al., 2020; Zhang et al., 2020a). The branch point of TDR/PXY and BAM/CLV1 clades is highly supported; these clades likely originated from gene duplication in a common ancestral gene before the divergence of vascular plants and bryophytes. Subsequently, TDR/PXY was lost in the lineage that includes *P. patens*. In the TDR/PXY branch, a gene duplication before the appearance of the angiosperms gave rise to a *PXY-LIKE* (*PXL*) gene, which was further duplicated into *PXL1* and *PXL2* in the eudicot lineage (Figure 2F). The BAM/CLV branch expanded through gene duplications, resulting in three *BAM* genes in addition to the *CLV1*gene in Arabidopsis (Figure 2G).

Our analyses indicate that the 27 *LRXI* Arabidopsis genes originate from six or seven genes (orthologues of *GSO* and/or *PEPR*, *At1g17230*, *RGI*, *HSL*, *TDR*, *BAM*) present in a common ancestor of vascular plants and bryophytes that have undergone duplications (Table 2).

*S. moellendorffii* has in total 14 genes with one to three paralogues for each founder gene. However, in the three bryophytes one or two of the founder genes have been lost, while in *A. trichopoda* orthologues of 18 of the Arabidopsis *LRXI* genes are present. Gene duplications followed by neofunctionalization with regards to ligand interaction may have refined cell-to-cell communication and facilitated the (r)evolution of flowering plants. Furthermore, nine of the present-day Arabidopsis *LRXIs* are likely results of genome duplication during the evolution of the eudicot species (Table 2) (Vanneste et al., 2014).

### LRRs and KDs evolve at different pace

Mutational changes in *LRXI* receptor genes may affect the output from the KD or the perception of peptide ligands through the ectodomain. The two parts of these receptors may therefore experience different selective pressure. We noted that for most of the clades in the tanglegram, the branches were longer in the LRR ectodomain tree compared to the KD tree (Figure 2). The branch length is proportional to the number of amino acid changes per site; therefore, this difference suggested that the LRR evolved faster than the KD. To further investigate the rate of evolution in *LRXI* genes of the 21 angiosperm subclades as well as six AMPS subclades, we computed the pairwise ratio (ω) between non-synonymous substitutions at non-synonymous sites (K_a_) and synonymous substitutions at synonymous sites (K_s_) for the DNA sequences encoding the KDs and the ectodomains (Figure 3). Comparison of the ratio between non-synonymous and synonymous substitutions is an effective method for detection of positive or purifying selection. In general, a ω of 1 indicates neutral evolution without any selective pressure, while ω > 1 indicates positive selection and ω < 1 purifying selection (Yang and Bielawski, 2000).

**Figure 3.**
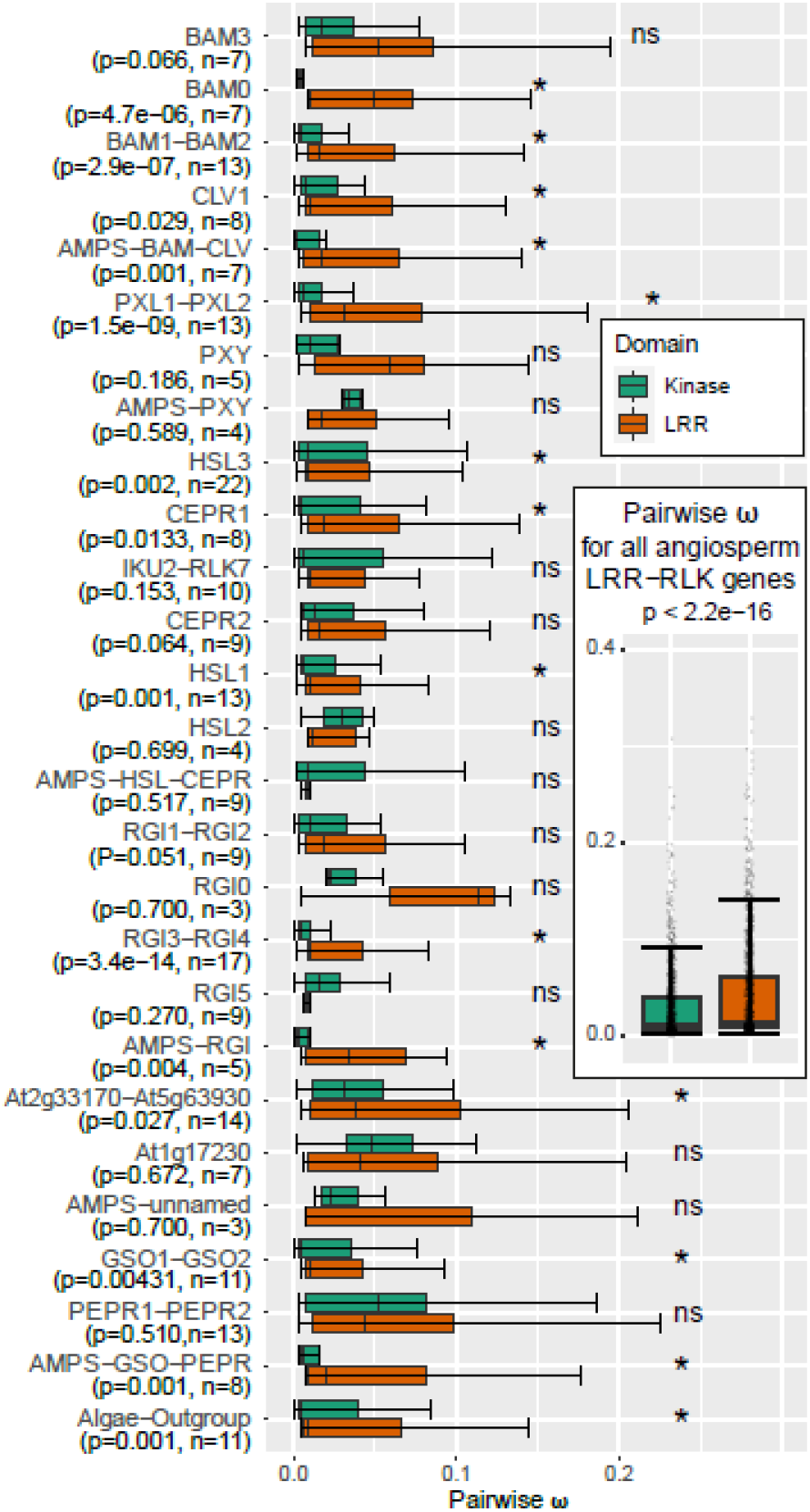
Ratio of nonsynonymous and synonymous substitutions in the angiosperm and AMPS LRXI gene sequences encoding the kinase domains (KDs) and LRR ectodomains. The seven clades were divided into subclades defined from the tanglegram in Figure 2 and detailed in Supplemental Figure 2, and the pairwise ratio (ω) between non-synonymous substitutions at non synonymous sites (Ka) and synonymous substitutions at synonymous sites (Ks) was calculated for each subclade. The boxplots delineate the mean and the interquartile range of ω. The error bars extend from a value 1.5 times smaller to a value 1.5 times larger than the interquartile range, p-values are based on a Wilcoxon test, n - number of sequences tested in the subclade. * - subclades with significant difference between ω_LRR_ and ω_KD_ (p < 0.05), “ns” - not significant (p> 0.05). Insert: ω ratio between all pairs of sequences, both within and between subclades. Each pairwise comparison is marked with a point. Outliers have been removed from all boxplots for clarity.

A recent survey of the evolution of *LRR-RLK* genes in angiosperm did not find the *LRXI* genes to be under significant positive selective pressure compared to other *LRR-RLK* genes (Fischer et al., 2016). We instead wanted to investigate if KDs of *LRXI* genes are under different selective pressure relative to the LRR domains. The ω ratios were therefore calculated separately for the two domains, and the Wilcoxon rank-sum test, which calculate the difference between sets of pairs to establish if these differences are statistically significantly different from one another, was used to calculate p-values. While the ω < 1 indicated purifying selection for both domains, the median ratio for all the angiosperm *LRXI* genes was significantly lower for *KD* (ω_*KD*_=0.015) than for the *LRR* (ω*_LRR_*=0.024, p-value < 2.2*10^−16^, Figure 3), supporting the idea that the KD is under a stronger purifying selective pressure than the LRR ectodomain.

This was a common trend for the angiosperm *LRXI* genes, as ω_*KD*_ of 15 of the 21 subclades was significantly lower than the ω_*LRR*_ according to the Wilcoxon test (significance level p<0.05), implying that the *LRRs* have evolved faster relative to the KDs.

Interestingly, the angiosperm subclades with the most significant differences between ω_*KD*_ and ω_*LRR*_ were enriched in duplicated monocot genes, and showed particularly low ω_*KD*_ values (Figure 3 and Table 3). Another interesting finding was the exceptionally low ω_*KD*_ for the BAM subclades (except for BAM3), where the receptors from bryophytes to Arabidopsis are known to be involved in control of meristem activity. The only subclade for which ω_*KD*_ > ω_*LRR*_ (p-value 0.002), was HSL3, while for the rest of the clades the difference was insignificant. The HSL3 subclade contains the highest number of sequences with several duplications in both monocots and dicots (Figure 2, and Supplemental Figure 1), suggesting that with the expanded number of genes the selective pressure is relaxed on the KD relative to the LRR. Among the AMPS subclades significant ω_*KD*_ < ω_*LRR*_ was found for the sister subclades to *RGI* (p-value 0.004, AMPS-RGI in Figure 3), and to GSO-PEPR (p-value 0.001), while there were no significant differences for the remaining subclades.

**Table 3.** Median values of pairwise K_a_/K_s_ (ω) for subclades of *LRXI* genes with p-values from Wilcoxcon rank-sum test.

| Subclade<sup>1</sup> | $\omega_{KD}$ | $\omega_{LRR}$ | p-value<sup>2</sup> |
| --- | --- | --- | --- |
| AMPS-GSO-PEPR | 0.0066 | 0.0203 | **0.001** |
| PEPR1-PEPR2 | 0.0531 | 0.0436 | 0.510 |
| GSO1-GSO2 | 0.0056 | 0.0112 | **0.004** |
| AMPS-unnamed | 0.0230 | 0.0081 | 0.700 |
| At1g17230 | 0.0490 | 0.0409 | 0.672 |
| At2g33170-At563930 | 0.0323 | 0.0386 | **0.027** |
| AMPS-RGI | 0.0033 | 0.0346 | **0.004** |
| RGI5 | 0.0166 | 0.0096 | 0.270 |
| RGI3-RGI4 | 0.0055 | 0.0111 | **3.4x10<sup>-14</sup>** |
| RGI0 | 0.0231 | 0.1142 | 0.700 |
| RGI1-RGI2 | 0.0106 | 0.0192 | 0.051 |
| AMPS-HSL-CEPR | 0.0090 | 0.0093 | 0.517 |
| HSL2 | 0.0298 | 0.0119 | 0.699 |
| HSL1 | 0.0060 | 0.0109 | **0.001** |
| CEPR2 | 0.0135 | 0.0171 | 0.064 |
| IKU2-RKL7 | 0.0070 | 0.0114 | 0.153 |
| CEPR1 | 0.0054 | 0.0187 | **0.013** |
| HSL3 | 0.0100 | 0.0099 | *0.002* |
| AMPS-PXY | 0.0351 | 0.0173 | 0.589 |
| PXY | 0.0207 | 0.0245 | 0.186 |
| PXL1-PXL2 | 0.0068 | 0.0313 | **1.5x10<sup>-09</sup>** |
| AMPS-BAM-CLV | 0.0029 | 0.0174 | **0.001** |
| CLV1 | 0.0083 | 0.0110 | **0.029** |
| BAM1-BAM2 | 0.0053 | 0.0169 | **2.9x10<sup>-07</sup>** |
| BAM0 | 0.0047 | 0.0492 | **4.7x10<sup>-05</sup>** |
| BAM3 | 0.0176 | 0.0523 | 0.066 |

<sup>1</sup>See Supplemental Figure 1 for subclade classifications.
<sup>2</sup>Significantly lower $\omega_{KD}$ than $\omega_{LRR}$ in bold ( $p < 0.05$ ), significantly higher $\omega_{KD}$ than $\omega_{LRR}$ in italics ( $p < 0.05$ ).

The observed difference in selection pressure is likely due to the differential structural and functional requirements facing the respective domains. The stronger conservation of the amino acids in the KD may reflect a need for preservation of an overall configuration to secure the kinase activity by maintaining a compact structure of alpha-helices and beta-sheets based on 13 tightly positioned conserved motifs (Liu et al., 2017). The ectodomain on the other hand needs a structural scaffold guaranteed by the repeated leucines, in addition to amino acid residues involved in peptide recognition. The necessity of molecular interaction between signaling peptides and their receptors must have set constraints on mutational changes of amino acids with properties and locations directly needed for ligand binding and receptor activation. However, as the peptide ligands vary in sequence and length, the LRRs must have sequence differences adapted to their specific ligand(s).

Additionally, the length of the whole ectodomain must fit the length of the peptide ligand, given that PTMPs interact along the inner surface of the ectodomain (Figure 4A-B). PEPRs and GSOs have for instance 26 and 31 LRRs, respectively, while most LRXI have 21-23 repeats. Correspondingly, the peptides interacting with PEPR and GSO are longer (17 and 20 amino acids, respectively) than the common 12-14 amino acids of the ligands of other subfamily XI receptors (Tang et al., 2015; Okuda et al., 2020). Thus, in searches for unidentified ligands, it is important to take into account the number of LRRs of the receptor of interest.

**Figure 4.**
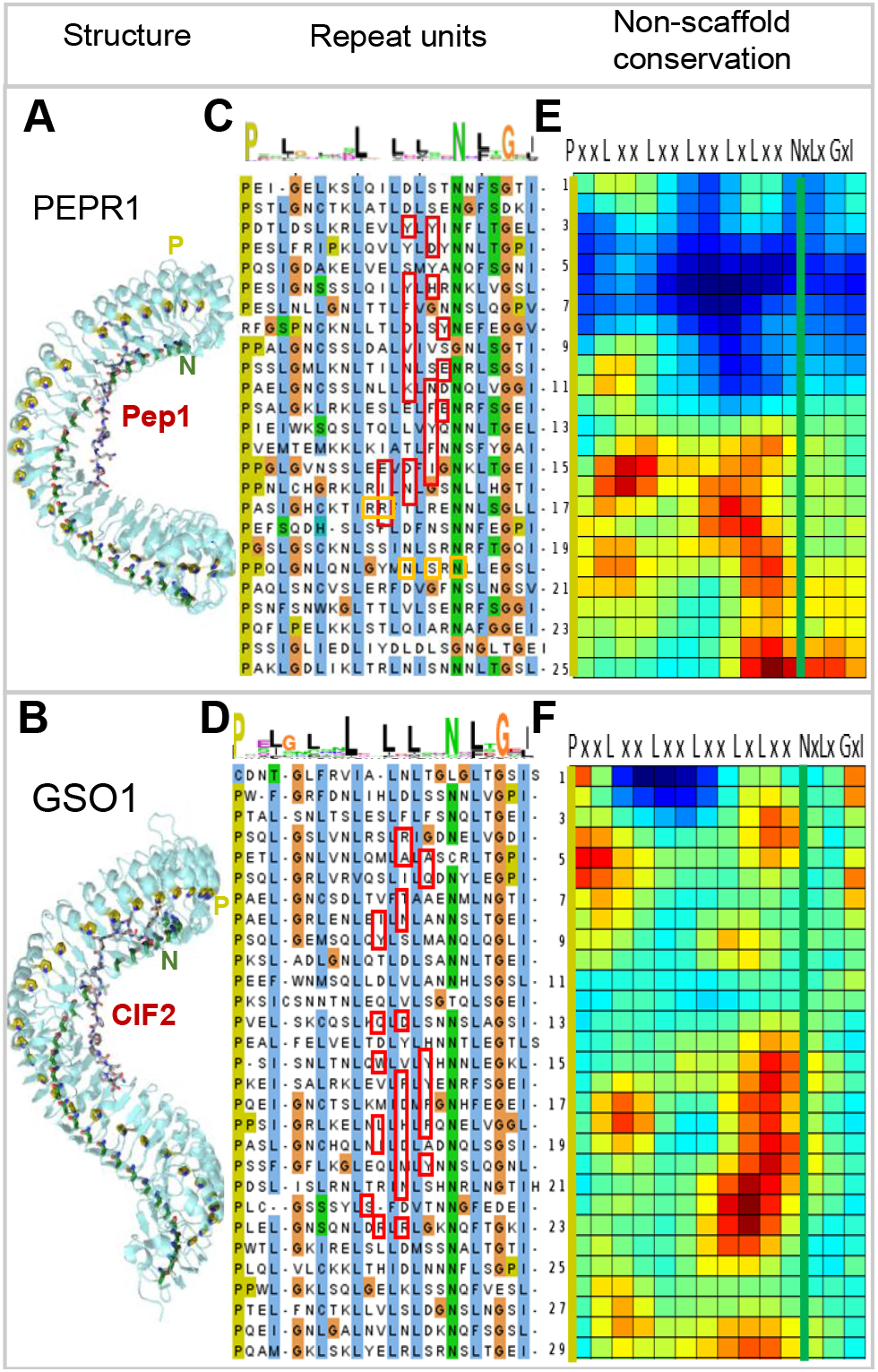
Conservation of LRRs scaffold residues and ligand-interacting amino acids. **(A)** and **(B)** Crystal structure of PEPR1 (PDBid:5GR8) with the ligand Pep1 (Tang et al., 2015) and GSO1 (PDBid: 6S6Q) with the ligand CIF2 (Okuda et al., 2020), respectively, interacting along the inner side of the LRR structures. The conserved Pro (P) and Asn (N) residues of the scaffolds are highlighted in mustard and green colors. **(C)** and **(D)** Alignment of the 24 amino acids long leucine-rich repeat units of PEPR1 and GSO1 with coloration based on amino acid properties. Above a WebLogo consensus sequence (http://weblogo.berkeley.edu/) visualizing the conservation of the scaffold residues. Residues of the LRR that according to the crystal structures shown in (A) and (D) interact with residues of the respective peptides, are marked with red rectangles, while orange squares represent residues interacting with the co-receptor BAK1/SERK3. **(E)** and **(F)** Heat maps generated using Repeat Conservation Mapping (http://www.bentlab.russell.wisc.edu/main/main.php) reflecting the degree of amino acid identify and similarity in a given position (X-axis) and a given repeat (Y-axis) for the non-scaffold residues of PEPR and GSO orthologues, respectively. The most conserved residues are in red and least conserved in blue. The position of the conserved Ps and Ns of the scaffolds are indicated by mustard and green colored vertical lines, respectively. Note that the conserved amino acid residues of PEPR orthologues (LRR 13 and onwards) are not coinciding with the majority of the amino acids of PEPR1 interacting with the Pep1 ligand (LRRs 2-13), but rather with the co-receptor-interacting residues. In contrast, ligand-interacting residues overlap with a substantial number of conserved residues in GSO orthologues (LRRs 14-23).

#### Methods for in silico homology mapping and modeling of peptide receptor interactions

Having identified presumptive orthologues of receptors and peptide ligands the challenge is to substantiate that candidates fit together as expected for a signaling module. *In silico* methods can be used to identify potential important residues in the LRRs. The simplest is to generate a visual representation of conserved of amino acids in alignments of putative receptor orthologues or members of peptide families, for instance using WebLogo (https://weblogo.berkeley.edu) (Crooks et al., 2004). Each position in the alignment has a stack of amino acid symbols and the height of the stack indicate the overall conservation at that position, while the relative height of the symbols at a given position indicates the relative frequency of specific residues (Crooks et al., 2004).

It may also be useful to generate a sequence logo of aligned repeats from single LRR ectodomain as that will demonstrate the scaffold amino acids as the most conserved residues in these repeats (see Figure 4C-D). The land plant LRR motif is 24 residues long and has specific amino acids, including six leucines, conserved in nearly fixed positions with the consensus sequence of PxxLxxLxxLxxLxLxxNxLxGxI (P – Pro; L – Leu; N – Asn: G – Gly; I – Ile; x – any amino acid). The LRR units that are most importantly involved in peptide binding are often deviating somewhat from the strict consensus of the scaffold (Figure 4 and Supplemental Figure 3). Conserved non-scaffold amino acids are on the other hand likely to represent functional sites, including residues interacting with ligands or co-receptors (Helft et al., 2011; Orr and Aalen, 2017).

Repeat Conservation Mapping (RCM) (http://www.bentlab.russell.wisc.edu/main/main.php) is a computational method that removes the scaffold residues from the alignment of orthologous LRRs and thereafter calculates the conservation of the remaining residues, presented as a heat map reflecting the degree of amino acid identify and similarity in a given position in a given repeat (Helft et al., 2011; Orr and Aalen, 2017). RCM can be used to identify conserved residues in orthologous receptors, which opens for *in vitro* and *in planta* experimental testing of functional importance.

The 3-dimentional (3D) structure of ectodomains bound to ligands are crucial for an understanding of the specificity of ligand interaction. The crystal structure of a number of Arabidopsis receptors and their peptide ligands have been solved over the last few years (Chakraborty et al., 2019) and RCM predictions have been validated by comparison to such crystal structures (Koller and Bent, 2014; Shi et al., 2019).

Solved crystal structures can also be used as templates in comparative modelling. 3D protein models can be generated by extrapolating coordinates of solved crystal structures of Arabidopsis peptide-receptor pairs to evolutionarily related peptides and receptors, using SWISS-MODEL (Arnold et al., 2006). Provided with the amino acid sequence of a target this program can search for suitable templates in the Protein Data Bank (PDB), which is growing rapidly. Lately SWISS-modeling has been extended to protein complexes (Waterhouse et al., 2018) which may be useful, since LRR-RLKs not only interacts with ligands, but also interact with other receptors and with co-receptors (Smakowska-Luzan et al., 2018; Chakraborty et al., 2019). Small co-receptors as key partners in signaling complexes (Ma et al., 2016; Hohmann et al., 2018) is a topic of its own right, which awaits to be elucidated in an evolutionary perspective in land plants.

### Conservation of amino acid residues in the ectodomain

When comparing LRR-RLKs one should keep in mind that high similarity observed between LRRs using BLASTp may result primarily from the conserved amino acids making up nearly half of each LRR. Thus, high BLASTp scores may not necessarily imply a conserved mode of ligand binding. One example is PEPR1 which in Arabidopsis is involved in amplification of biotic and abiotic stress responses and interact with endogenous stress-induced Pep peptides (Safaeizadeh and Boller, 2019). The phylogenetic distribution of the Pep peptides is, however, limited to angiosperms, and even within flowering plants Pep peptides show interfamily incompatibility. Peps of heterologous origin cannot be recognized as ligands due to rapid co-evolution of PEPR LRRs with PEPs (Lori et al., 2015). Consistent with this the ω_*LRR*_ of PEPR1-PEPR2 subclade has a higher value than most of the LRXI subclades (Table 3). The scaffold residues as well as putative co-receptor-interacting residues are conserved, but the majority of the Arabidopsis PEPR1 residues interacting with the peptide ligand Pep (according to the solved crystal structure (Tang et al., 2015)), are not conserved among PEPR orthologues from the species used in our studies (Figure 4A, C and E).

This situation has likewise been found for an LRR-RLK of subfamily XII involved in defense, the FLAGELLIN-SENSITIVE2 (FLS2) which, like PEPR, is interacting with the co-receptor BAK1/SERK3 (Koller and Bent, 2014). This is in contrast to the conserved ligand-interacting residues of receptors involved in developmental processes, like GSO1 interacting with the confirmed ligand CIF2 (Figure 4B and F) (Okuda et al., 2020) or the *M. polymorpha* and *S. moellendorffii* orthologues of HSL2 interacting with putative IDA peptides of these species (Supplemental Figure 3).

### Matching DY^sulf^ peptides with receptors

An additional twist when investigating potential peptide-receptor interactions is the presence of modified amino acid residues, in particular sulfated Tyr and hydroxylated Pro. We have confirmed that GSO and RGI homologues are present in non-flowering plants (Bowman et al., 2017; Liu et al., 2017; Man et al., 2020), and hypothesized that their respective ligands are conserved in the species where presumptive receptors are present. Using PHI-BLASTp, as described, we indeed retrieved ligand candidates from several species (Figure 1 and 5). The candidates show limited sequence identity (Figure 5A), however this is also the case for the four Arabidopsis CIF peptides and the recently identified TWISTED SEED1 (TWS1) peptide which show differential affinities to GSO1 and GSO2 (Barbosa et al., 2019) (Fiume et al., 2016; Doll et al., 2020; Okuda et al., 2020). TWS1 functions with the GSOs to form a functional cuticle around the developing embryo of Arabidopsis.(Tsuwamoto et al., 2008).

**Figure 5.**
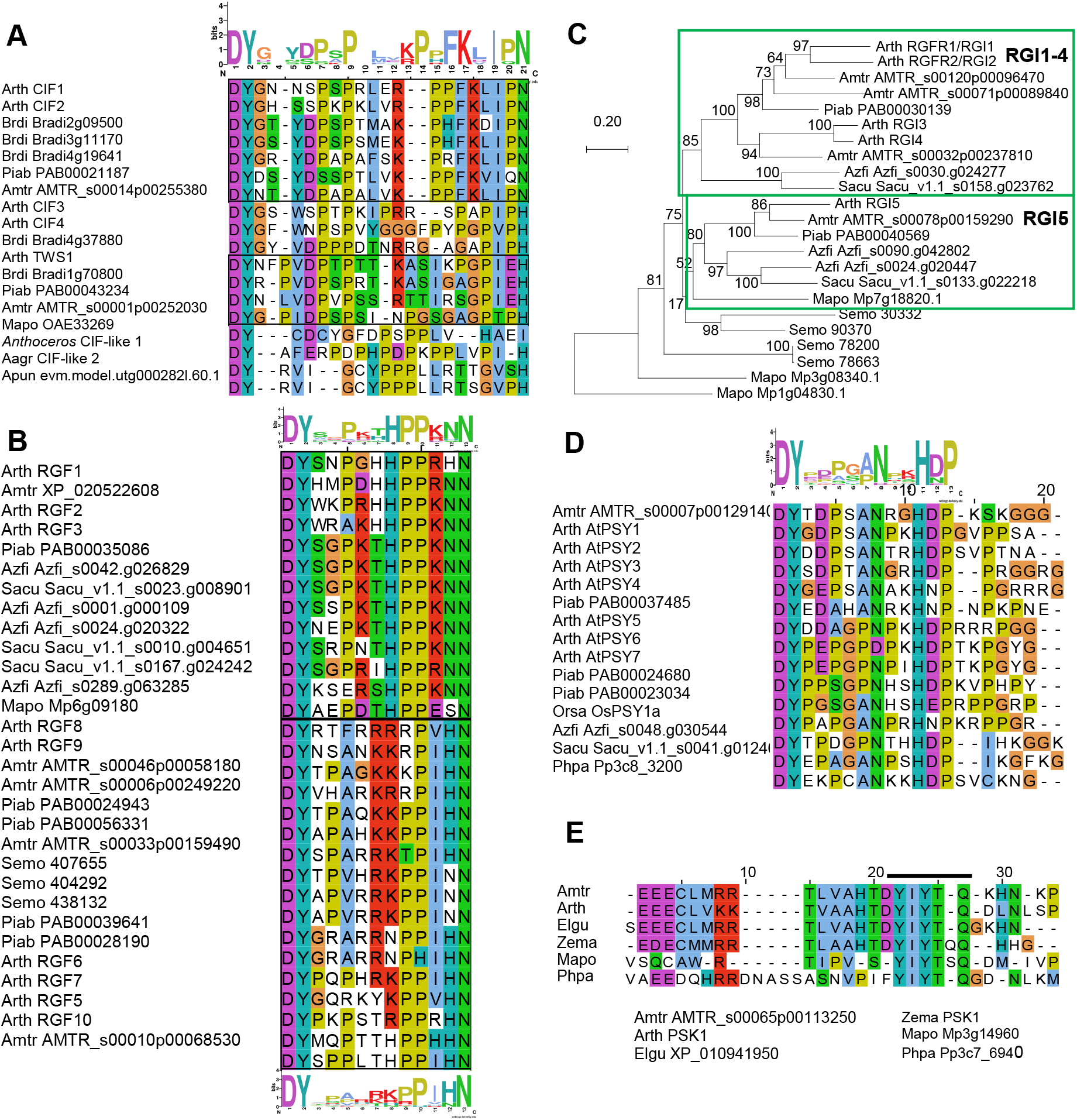
Conservation of sulfated peptides. **(A)** Alignment of CIF- and TWS1-related peptides. Above the alignment shown is the sequence logo of the top seven sequences (boxed), which represent canonical CIFs. TWS1 and its putative orthologues are also boxed. **(B)** Alignment of the presumptive RGF peptides, with amino acids colored according to chemical properties. RGFs are classified into two distinct groups (boxed), represented by the sequence logos above and below the alignment. **(C)** Phylogenetic tree based on the ectodomain amino acid sequences of RGIs in Arabidopsis, and their orthologues in the *A. trichopoda,* the gymnosperm *P. abies*, the ferns *A. filiculoides* and *S. cucullata,* the lycophyte *S. moellendorffii* and the bryophyte *M. polymorpha* constructed by the maximum likelihood method. The GSO homologue of *M. polymorpha* (Mp1g04830.1) from the neighboring clade was used as outgroup. Bootstrap values were obtained from 500 bootstrap replicates and indicated at the nodes as percentages. The tree is drawn to scale, with branch lengths measured in the number of substitutions per site. **(D)** Alignment of the presumptive PSY peptides, with a sequence logo of the first 13 amino acids above the alignment. **(E)** Alignment of the PSK1 mature peptide (YIYTQ) and adjacent conserved motif. Sequence logos were generated using WebLogo 2.8.2 (https://weblogo.berkeley.edu).

It may seem difficult to tell CIF and RGF peptides apart since they both start with DY (where Y is sulfated), they are enriched in Pro residues, and often end with a C-terminal Asn (N). They do, however, differ in length (Figure 5A-B, Table 1). The solved crystal structure of RGF1 and its receptor RGI3 (At4g26540) (Song et al., 2016); (note that the *RGFR/RGI* gene nomenclature differs in the literature) revealed that the DY^sulf^ residues are recognized by the amino acid motif RxGG of the receptor. Importantly, the RGI orthologues from different land plant lineages contain this motif, which distinguishes RGIs from other LRXI members (Supplemental Figure 4).

To further clarify phylogenetic relationships among RGI homologues from different lineages, we bridged the evolutionary gap between Arabidopsis and *S. moellendorffii* by finding genes encoding highly similar receptors in ferns (duckweed, *Azolla filiculoide*, and buce plant, *Salvinia cucullata*) and gymnosperm (Norway spruce, *Picea abies*). In each of these species, two or three *RGI* genes were identified, one or two belonging to a clade that includes Arabidopsis *RGI5* and the other to a branch with three *A. trichopoda* and four Arabidopsis *RGIs* genes (Figure 5C), suggesting one gene duplication in the common ancestor ferns and gymnosperms generated the RGI1-4 and RGI5 clades. In the RGI1-4 clade, at least three gene duplications are inferred: one in the seed plant lineage, generating two subgroups, both of which experienced another duplication in the eudicot lineage. A previous study also indicates that gene copies increased through lineage-specific gene duplications (Man et al., 2020).

In Arabidopsis, Tyr sulfation regulates protein-protein interactions and affects receptor binding (Matsubayashi and Sakagami, 1996; Shinohara et al., 2016; Kaufmann and Sauter, 2019). The gene encoding the responsible enzyme, *TYROSYLPROTEIN SULFOTRANSFERASE* (*AtTPST*), was discovered in Arabidopsis (Komori et al., 2009). TPST-encoding genes are conserved across land plants and found in *S. moellendorffii*, *P. patens*, and *M. polymorpha*, as well in ferns and gymnosperms (Supplemental Figure 5) supporting the likely presence of sulfated signaling peptides and the plausible contribution of PTMs to the ligand diversity in early land plant evolution.

PSY1 and phytosulfokine (PSK) are two sulfated peptides of 18 and five amino acid, respectively. PSY1 has an N-terminal DY^sulf^, and promotes cellular proliferation and expansion (Amano et al., 2007). The mature PSK peptide is only five amino acids long (YIYIQ) and promotes cell growth, acts in the root apical meristem, contributes to pollen tube guidance, and integrates growth and defense (Sauter, 2015; Kaufmann and Sauter, 2019). We identified putative orthologues both for PSY1 and PSK in *P. patens, M. polymorpha* as well as *P. abies* (Figure 5D-E). *PSK-*like genes and bioactivities have previously been detected in gymnosperms (Igasaki et al., 2003; Wu et al., 2019), but surprisingly, we did not find any orthologue of the Arabidopsis PSY1-receptror in the bryophytes. In contrast to all the subfamily XI receptors discussed here, the two PSK-RECEPTORs (PSKRs) of Arabidopsis belong to the subfamily X, of which binding to small ligands is facilitated by a so-called island domain (ID) in the LRR structure that doesn’t fit in with the rest of the repeats. In the PSK-like sequences found in *P. patens* and *M. polymorpha* the Asp (D) residue preceding the YIYIQ sequence is missing, but a conserved N-terminal region can still be recognized (Figure 5E). Such conserved sequences outside the assumed mature peptide might be involved in precursor processing and peptide maturation steps, which have been less characterized in plant peptidic signals.

### Matching receptors and hydroxylated ligands

*In silico* searches for homologues of many known Arabidopsis peptide sequences have identified candidate ligand-receptor pairs across land plants, with the latest discovery of BAM/CLV1 receptor and an H-type CLE peptide in hornworts (Li et al., 2020; Zhang et al., 2020a). This was further complemented by our findings of hornwort TDR/PXY and R-type CLE homologs (Figures 1; Table 2). It is, therefore, feasible that both H-CLE and R-CLE signaling pathways are conserved in hornworts.

Putative orthologues of IDA and HSL2 were also identified in *M. polymorpha* (Bowman et al., 2017) and therefore we looked for IDA in other AMPS. We did not detect IDA orthologues in the moss, but identified potential hits in *S. moellendorffii* and *Anthoceros* hornworts (Figure1 and 6). The crystal structure of synthetic AtIDA peptide bound to its receptor AtHAE has previously been solved, and showed as expected that the ligand is positioned along the inner surface of the LRR ectodomain of the receptor (Figure 6A) (Santiago et al., 2016). Using this structure (PDBid:5ixq), we have recently generated a three-dimensional model of IDA-HSL2 orthologues from oil palm, thereby substantiating a likely interaction in this monocot species (Shi et al., 2019). Here we have used the same modeling strategy for putative peptide ligands and receptors of *M. polymorpha* and *S. moellendorffii*, which demonstrates that the potential overall hydrogen bonding interactions between the modeled MpHSL2 and SmHSL2 receptors and their corresponding IDA peptides (MpIDA1-2 and SmIDA1-3) support receptor-peptide binding, regardless of variations in IDA-peptide sequences (Figure 6B-C). A substantial fraction of the AtHAE amino acids generating hydrogen bonds with IDA residues are identical in MpHSL2 and SmHSL2 (Figure 6A-B and Supplemental Figure 3), and crucial residues, in particular the central Pro and the C-terminal His-Asn, are also conserved in the amino acid sequence of the MpIDA1-2 and SmIDA peptides (Figure 6C). Thus, the generated models are consistent with the possible function as ligand-receptor pairs in early-diverging land plant lineages.

**Figure 6.**
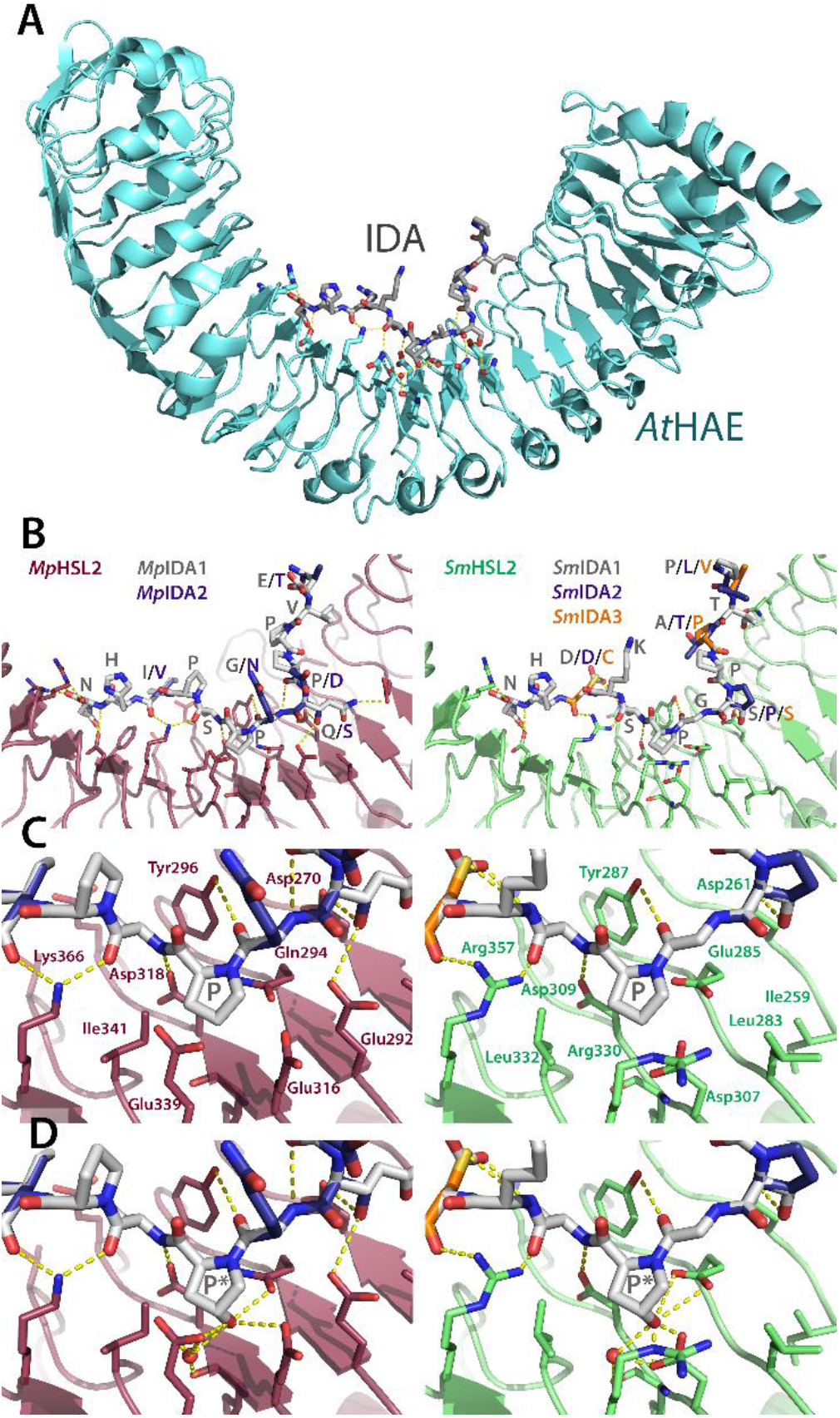
Modelling of the interaction between putative IDA ligands and HSL2 receptors of *M. polymorpha* and *S. moellendorffii.* **(A)** The AtHAE-AtIDA crystal structure (PDBid:5ixq) (Santiago et al., 2016) the IDA peptide is lining up along the inner face of the LRR structure. **(B)** Overall models of the interaction between *M. polymorpha* and *S. moellendorffii* putative HSL2 receptors (MpHSL2 in magenta and SmHSL2 in green) with respectively two and three *M. polymorpha* and *S. moellendorffii* superimposed putative IDA peptides (grey backbones) built on the AtHAE-AtIDA structure using SWISS-MODEL (Arnold et al., 2006). Note in particular receptor interaction with the Asn (N) at the C-terminal end of the peptides. **(C)** Close-up view of the central parts of the respective receptor models and the surrounding hydrogen bonding network, with a central Pro (P) in the ligands. **(D)** Close-up view as in (C), however, with hydroxylation of the central Pro (P*), which facilitates formation of additional hydrogen bonds. Central amino acids of the receptors, as well as the peptides, are shown as sticks and colored by atom type. Water molecules are shown as red spheres, and modelled based on coordinates from the AtHAE-AtIDA crystal structure. Hydrogen bonds are depicted as dotted lines (yellow). Residues involved in hydrogen bonding to the peptides are depicted with three-letter symbols in colors according to the respective structures. The peptide residues are shown in one-letter symbols. All structure figures were prepared using PyMOL (Schrödinger, LLC).

Synthetic peptides of the IDA/IDL family have been shown to bind and activate the HAE and HSL2 receptors more efficiently when the central Pro is hydroxylated (Table 1) (Butenko et al., 2014; Santiago et al., 2016), and this PTM has also been detected in planta on CLE peptides (Stührwohldt and Schaller, 2019). We therefore modeled MpHSL2 and SmHSL2 with hydroxylated MpIDA and SmIDA peptides. In these models, the hydroxylation postulated an increased number of hydrogen bonds, which would result in a stronger interaction (Figure 6C and D). Interestingly, genes encoding prolyl 4-hydroxylases (P4Hs) which mediate Pro hydroxylation, are evolutionarily conserved from algae to angiosperms (Myllyharju, 2003; Ogawa-Ohnishi et al., 2013). Thus, mature peptides produced from IDA and CLE orthologues may have been hydroxylated already in early land plants (Oelkers et al., 2008; Bowman et al., 2017). Hydroxyproline has also been found in members of other peptide families (Table 1).

In Arabidopsis, IDA and its receptors HAE and HSL2 are involved in cell separation processes, like floral organ abscission, lateral root emergence, and root cap sloughing (Aalen et al., 2013; Shi et al., 2018). In various angiosperm species that shed other organs, like fruits or leaves, the IDA-HAE/HSL2 signaling module has also been found expressed at the base of the organ to be shed (Shi et al., 2019), indicating that the molecular process underlying cell separation is highly conserved. This suggests that a change in timing or location of expression of peptide signaling components through changes in their cis-regulatory elements may have served as an important driving force for evolutionary changes in plant architecture and reproductive strategies.

Cell separation processes also play a role in bryophyte development, for example in air chamber formation of *M. polymorpha*. Ishizaki et al. (2015) revealed that E3 ubiquitin ligase regulates cell separation in *Marchantia* air chamber formation, and it is not known whether the IDL-HAE/HSL signaling module is involved or not in this process. It will be important to find out more about biological processes in bryophytes and lycophytes that are controlled by cell separation, where the putative IDA and HSL orthologues might have conserved roles.

### Evolution of peptide ligands

In the 20 years since Shiu and Bleecker identified the LRR-RLKs of Arabidopsis, the knowledge about their function and their confirmed ligands have gradually increased. Principally it is expected that genes encoding a receptor and its ligand display the same mutant phenotype. The CLAVATA1 (CLV1) receptor and the CLV3 peptide ligand were discovered already at the end of the last century by their spectacular meristem phenotypes (Clark et al., 1995; Clark et al., 1997). A double mutant of the closely related receptors HAE and HSL2 was needed to disclose their involvement in floral organ abscission, like their peptide ligand IDA (Butenko et al., 2003; Cho et al., 2008; Stenvik et al., 2008). Furthermore, triple to quintuple mutants were necessary to disclose the involvement of the RGI receptors in root development, with RGFs peptides as ligands (Shinohara et al., 2016; Song et al., 2016; Qiu et al., 2020). Thus, functional redundancy on the receptor side as well as the peptide side has made it complicated to identify ligand-receptor pairs through genetics. However, in non-seed plants with often fewer duplicated genes and with the establishment of CRISPR-Cas mutagenesis, a genetic approach for identification of peptide ligand receptor-pair should be more feasible. In order to pursue this, we need to develop more non-seed, genetically tractable model plants (Rensing, 2017).

It has just started to be revealed that the CLE peptide signaling has evolutionarily conserved roles from *P. patens* and *M. polymorpha* to angiosperms. One of the known key roles of the CLE signaling in angiosperms is to regulate cell proliferation in the sporophytic (2*n*) meristems, which grow indeterminately. In the bryophytes, the sporophyte shows determinate growth while indeterminate meristems are present in the gametophyte body (1*n*). The CLE signaling in both *P. patens* and *M. polymorpha* is involved in cell proliferation in the gametophyte and determination of the orientation of cell division planes which thereby facilitating the transition from 2D to 3D growth (Whitewoods et al., 2018; Hirakawa et al., 2019). Similarly, Arabidopsis quadruple mutant for *CLV1* and the three *BAM* genes show abnormal cell division planes. These studies support the idea that peptide-receptor modules are conserved and employed in a parallel manner, although the evolutionary relationship between the bryophyte gametophytic meristems and the vascular plant sporophytic meristems is still debated (Bowman, 2013).

To our surprise, several Arabidopsis CLE peptides have been found to interact with LRRs of different clades. The AtCLE9/CLE10 peptides interact with both the HSL1 and BAM1 receptors, depending on tissue type (Qian et al., 2018). AtCLE14 peptide can signal through AtPEPR2 (Gutierrez-Alanis et al., 2017). While PEPs and CLEs differ in length, the C-terminal 12 amino acids of mature PEPs share sequence similarity to CLEs (Table 1), offering a possible explanation for the recognition mechanism. Another CLE peptide, AtCLE45, which differs from other CLE peptides by a number of arginine (Arg, R) residues near its N-terminus, is perceived by canonical CLE receptors as well as an RGF receptor homolog, AtRGI4/SKM2 (Endo et al., 2013; Hazak et al., 2017).

These “unconventional” ligand-receptor pairs raise the question of how the specificity of ligand-receptor pairs evolves. It is of note that genes encoding CLE peptides highly similar to AtCLE9/CLE10 are present in both *M. polymorpha* and *P. patens* (PpCLE5/CLE6) (Whitewoods et al., 2018) (Figure 7). Our phylogenetic investigations suggest that founder receptors, representing the seven clades, by the onset of the angiosperm era had increased about three times, and in each of the clades a new gene has evolved during dicot evolution. We therefore speculate that during early evolution of peptide signaling, there were fewer peptides, and receptors were less specific. Accordingly, large-scale clustering analyses of land plant CLE peptides based on their entire prepropeptide sequences found a smaller number of clusters in AMPS, indicative of the diversification of CLE peptide sequences, which could have resulted in changes in the specificity of ligand-receptor interactions during land plant evolution (Goad et al., 2017). Hirakawa et al. (2017) found that a synthetic CLE peptide, KIN named after the K (2^nd^), I (10^th^), and N (12^th^) residues crucial for activity, exerts both R-type and H-type CLE activities and interact directly with both the CLV1 and the TDR receptors (Figure 7) (Hirakawa et al., 2017), illustrating the potential for a broader specificity in ligand-receptor interaction hidden in short mature peptide sequences (Figure 7).

**Figure 7.**
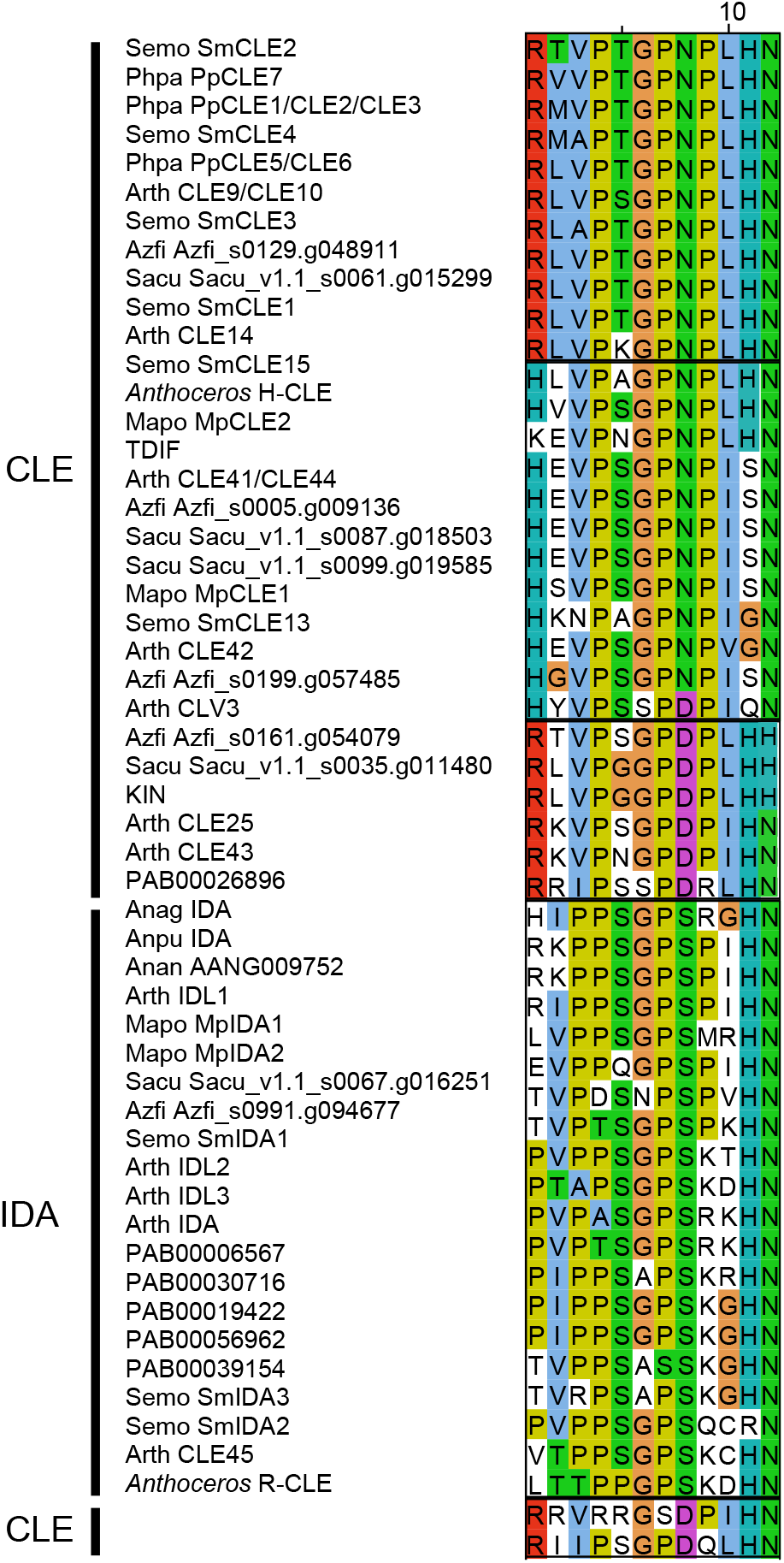
Conservation of CLE and IDA peptides. Alignments of mature CLE and IDA peptides in land plants and a synthetic peptide, KIN. Note shared core residues (PSGP) and C-terminal end (HN) between IDA and some of the CLE peptides, and the presence of almost identical R-CLE, H-CLE and IDA peptides in AMPS, ferns and Arabidopsis. The amino acids are colored according to chemical properties.

We may also learn from some common structural features in known endogenous peptide ligands (reviewed and discussed in (Zhang et al., 2016)). For instance, the IDA and CLE peptides are all 12-14 amino acids long, proline-rich, and need C-terminal HN or HH residues for function (Figure 7; Table 1). Similarities are also recognized among CEPs, PIP-PIPLs, and IDA peptides with respect to the size and amino acid composition. Most of the CEPs and PIP-PIPLs, however, lack the C-terminal HN or HH residues that have been shown to interact with two closely positioned arginine (Arg, R) residues found in many receptors (Hou et al., 2014; Vie et al., 2015; Song et al., 2016) (Table 1). Instead, CEPs and PIP-PIPLs share the C-terminal GxGH motif (Vie et al., 2015) (Table 1). The CEPs and PIP-PIPLs have been demonstrated to signal through closely-related receptors of the HSL major clade, CEPRs and RLK7, respectively (Figure 2). This points to the possibility that a ligand recognition mechanism is conserved among recently diversified receptors, and in the case of the CEP and PIP-PIPL signaling, the GxGH motif could be crucial for peptide-receptor interactions.

### Perspectives

With the increasing availability of genomic data, it is now possible to analyze the molecular inventory of peptide-signaling components in all major lineages of land plants. All known signaling peptides were originally discovered in angiosperms, however, the identification of LRXI in non-seed plants and our successful hunt for their likely peptide ligands, give a new perspective on the roles and evolution of cell-to-cell signaling. A fourth of the present-day XI receptors of Arabidopsis, with seemingly angiosperm-specific functions, have a history all the way back to the ancestor of the bryophytes and vascular plants. In addition to the previously identified orthologues of CLE peptide ligands and their receptors (BAM/CLV and TDR) (Whitewoods et al., 2018; Hirakawa et al., 2019; Hirakawa et al., 2020), our approach has disclosed early occurrence of orthologues of GSO and RGI receptors and their CIF and RGF ligands. Additionally, the direct interaction between HSL2 and IDA orthologues of *M. polymorpha* and *S. moellendorffii*, was substantiated by taking advantage of the solved 3D structure of the Arabidopsis peptide-receptor pair. We have also bridged the gap between AMPS and Arabidopsis by identifying likely orthologues of LRXI in ferns and gymnosperms. This may be a starting point for investigation of peptide signaling in these taxa. This would elucidate LRXI evolution prior to the emergence of angiosperms, and possibly identify common ancestors for receptors present both in *A. trichopoda* and Arabidopsis (Table 2). Nine of the Arabidopsis XI genes are results of gene duplication within the eudicots lineage (Table 2). Several of the receptor families have also proliferated in monocot lineages (Figure 2). This may reflect the increased need for cell-to-cell communication for development of more complex organisms with more complex organs. An approach similar to what we have shown here with the integration of phylogenetic, functional and structural information, should facilitate the disentanglement of distinct evolutionary histories of peptide ligands and their receptors in mono- and dicots. One study along these lines have recently been published on the BAM/CLV1/TDR receptors and CLE peptides (Cammarata and Scanlon, 2020).

Interestingly, the GSO/RGI clades on one side and the HSL/TDR/BAM/CLV clades on the other, seem to have ligands representing two different groups – those with sulfated Tyr and the CLE/IDA-like peptides. Founder genes for both were possibly present in the most recent common ancestor of vascular plants and bryophytes. Alternatively, convergent evolution cannot be completely dismissed and might account for the highly variable central part of peptide precursor sequences. Signaling peptides can evolve not only from changes in the primary sequences, but through changes in proteolytic processing of prepropeptides, or by PTM enzymes conferring Tyr sulfation or Pro hydroxylation. The PTM processes tend to be irreversible and hence require precise regulation, pointing to the possible contribution of PTMs in diversification of peptide ligands and in defining the specificity of ligand-receptor interaction since the early stages of land plant evolution. Our structure-based modeling of IDA-HSL2 is consistent with an ancient origin of peptide hydroxylation. Structural information can be used to model whether suggested peptide-receptor pairs actually fit together, and structural analyses of bioactive forms of PTMPs in AMPS will be critical for confirming the prediction from the models.

It needs to be mentioned that a number of receptors, even some in Arabidopsis, are orphans, in the sense that their peptide ligands are unknown. Likewise, all PTMP families seem to have orphan members. These may have resulted from gene or genome duplications where some sister genes have accumulated mutations that weakened ligand-receptor interaction. An alternative (and likely) explanation is that more specific genetic, molecular and biochemical factors and conditions for peptide-mediated cell-to-cell communication remain to be discovered. In line with this suggestion, CLE9/10 of Arabidopsis is recruited to bind HSL1 very efficiently in the presence of the small multi-faceted co-receptors SERK1-2 or 3 (Kd = 1.5 μM), while in the absence of SERK the preferred receptor is BAM1 (Kd = 0.112 μM) (Qian et al., 2018). To dig deeper into the evolution of peptide signaling, we recommend the matching of additional phylogenetic, functional and structural data, especially on the interaction with other XI receptors (Smakowska-Luzan et al., 2018), and co-receptors like the SERKs as key partners in receptor complexes in diverse peptide signaling pathways (He et al., 2018; Liang and Zhou, 2018; Hohmann et al., 2017; Ma et al., 2016).

Land plant evolution must successfully have made use of a broadened spectrum of receptors as more complex organ evolved and interspecies interactions increased. However, except for the bryophyte CLE signaling pathways, biological roles have not been assigned for peptide LRXI signaling other than in angiosperms. Peptide signaling pathways have most often been studied in angiosperm-specific organs, of which homologous structures are not universally found outside the flowering plants. A question therefore arises as to whether these peptide signaling modules mainly have undergone neofunctionalization in more recently evolved lineages. This may well be the case from an organismal perspective, however, from a molecular and cellular perspective it may not. Where and when a signaling system triggers a given outcome (e.g. cell division, cuticle formation or cell separation) may actually be the crux of the matter. Thus, a very important next step in tracing the evolutionary origin and history of land plant peptide signaling pathways will be to investigate thoroughly the temporal and spatial expression patterns of LRXI receptors and their putative peptide ligands, and to study downstream molecular events.

## Supplemental material

**Supplemental Table 1.** Major CRP families in land plants. Support for Table 1.

**Supplemental Table 2.** References for the signaling peptides. Support for Table 1.

**Supplemental Figure 1.** Phylogeny for subfamily XI LRR-RLKs based on the kinase domain and the LRR ectodomain. Support for Figure 2.

**Supplemental Figure 2.** Differential sequence similarities in the ectodomains and the kinase domains of CEPR1, CEPR2, RLK7 and IKU2 compared to HAE. Support for Figure 2.

**Supplemental Figure 3.** Conservation of amino acids residues involved in ligand binding. Support for Figure 4 and 6.

**Supplemental Figure 4.** Conservation of ligand-binding residues in RGI receptors. Support for Figure 5.

**Supplemental Figure 5.** Phylogenetic analysis of TYROSYLPROTEIN SULFOTRANSFERASE (TPST). Support for Figure 5.

**Supplemental Data Set 1.** Names and IDs for proteins and genes presented in Figures 1-6. Excel format.

**Supplemental Data Set 2.** Alignment of LRRs for phylogenetic analyses.

**Supplemental Data Set 3.** Alignment of kinase domains for phylogenetic analyses.

**Supplemental Data Set 4.** Phylogenetic tre file for LRRs.

**Supplemental Data Set 5.** Phylogenetic tre file for kinase domains.

## Acknowledgements

We apologize to authors of literature not discussed due to space constraints. We thank three anonymous reviewers for helpful comments that improved the manuscript. Investigations by CF and SS were supported by Japan Society for the Promotion of Science London (KAKENHI 18H05487, 20H00422, and 20KK0135 to SS and 20K06770 to CF) and research grants from the Nakatsuji Foresight Foundation and the Sumitomo Foundation to CF.

## Author Contributions

MW and SS initiated and drafted the work; CF, AKK, RMA and RBA identified and analyzed sequences; MH performed structure analyses and modelling; CF and RBA wrote the article. CF created Figure 1, 5 and 7, and Table 1 with supporting tables and figures with input from RMA and RBA; AKK generated Figures 2 and 3, and Table 2 and 3 with supporting figures with input from RMA and RBA; Figures 4 and 6 with supporting figures were created by MH and RBA.

## Distribution of materials

The authors responsible for distribution of materials integral to the findings presented in this article in accordance with the policy described in the Instructions for Authors (www.plantcell.org) are Chihiro Furumizu and Reidunn B. Aalen.

