## Supplemental Figure 1 for "The Sequenced Genomes of Non-Flowering Land Plants Reveal the (R)Evolutionary History of Peptide Signaling"

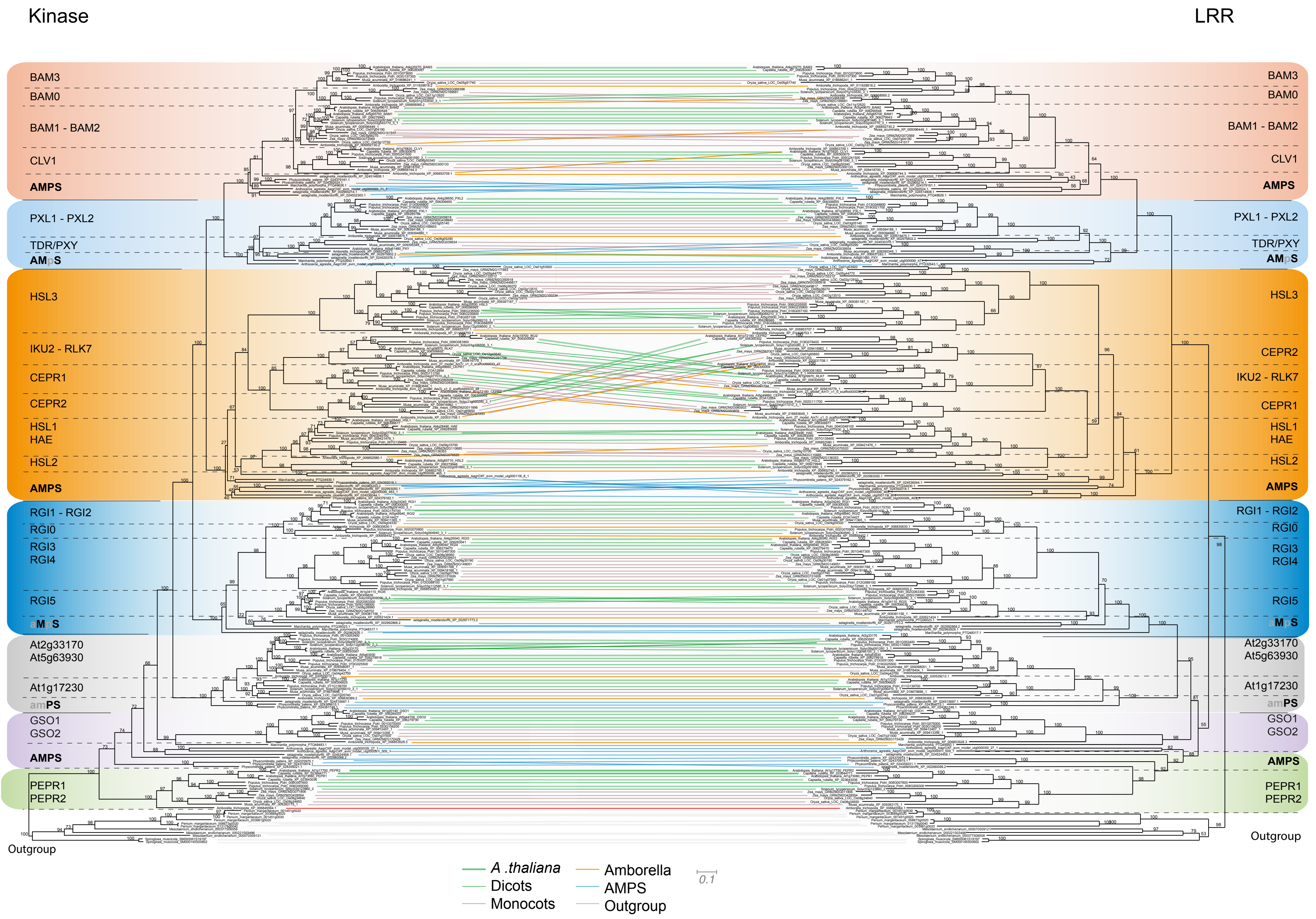


**Supplemental Figure 1. Tanglegram of the kinase domains (KDs) and the LRR ectodomains encoded by LRXI genes. Supplement to Figure 2.**

The phylogenetic trees were constructed separately for the LRRs and the KDs. A tanglegram connects KD and LRR originating from the same LRXI. The connecting lines represents *A. thaliana* (dark green); the dicotsc *Capsella rubell*, *Populus trichocarpa* and *Solanum lysopersicum* species (light green); the monocots *Musa acuminata*, *Oryza sativa*, and *Zea mays* (brown); the basal angiosperm *A. trichopoda* (orange with boxes); the hornwort *Anthoceros*, the liverwort *M. polymorpha*, the moss *P. patens*, and the lycophyte *S. moellendorffii* (blue, collectively referred to as AMPS), and three species of Zygnematophyceae algae used as outgroup (grey). Connecting lines between monophyletic groups of the same kind (i.e. either monocots, dicots, *A. trichopoda*, AMPS, or the outgroup) have been collapsed for clarity. Branches drawn with solid lines are well supported (SH-aLRT > 0.85), while dashed lines represent unsupported branches with SH-aLRT < 0.85. Support values for all branches can be found in Supplemental Figure 2. The scale bar for branch lengths represents the average number of changes in amino acids per site, and the two trees of the tanglegram are drawn to scale. The seven colored boxes (A)-(G) represent monophyletic clades of the indicated angiosperm LRXIs with a common origin and a last common ancestor with one or more AMPS (see Table 2 for details). The braches representing Arabidopsis KDs or LRRs are labeled at the tips.
