## Supplemental Figure 2-5 for "The Sequenced Genomes of Non-Flowering Land Plants Reveal the (R)Evolutionary History of Peptide Signaling"

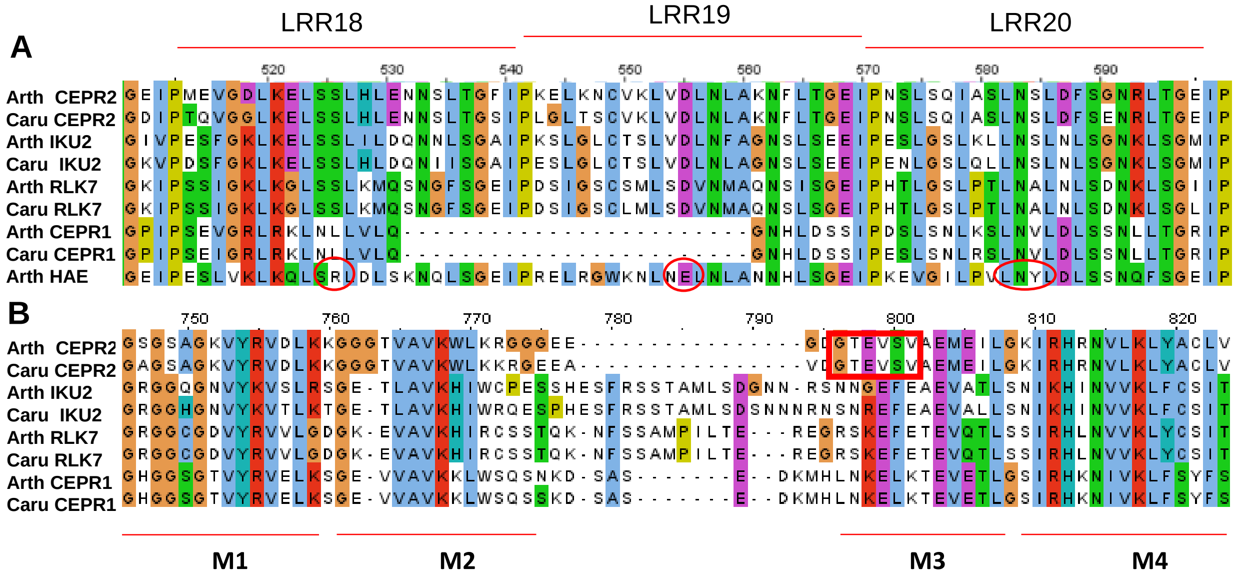


**Supplemental Figure 2.** Differential sequence similarities in the ectodomains and the kinase domains of CEPR1, CEPR2, RLK7 and IKU2 compared to HAE. Support for Figure 2.

**(A)** Alignment of the three repeats of the ectodomain of the indicated receptors from *A. thaliana* and *Capsella rubella* corresponding to the Arabidopsis HAE retreats LRR18, LRR19 and LRR20. Amino acids corresponding to the HAE residues interacting with the co-receptor SERK1 are encircled (Santiago et al. 2016).

**(B)** Alignment of the N-terminal part the kinase domain of the indicated receptors from *A. thaliana* and *Capsella rubella.* The first four conserved motives of the kinase domain (Liu et al. 2017) are indicated below the alignment. The deviating M3 motif of CEPR2 is indicated by a red rectangle.

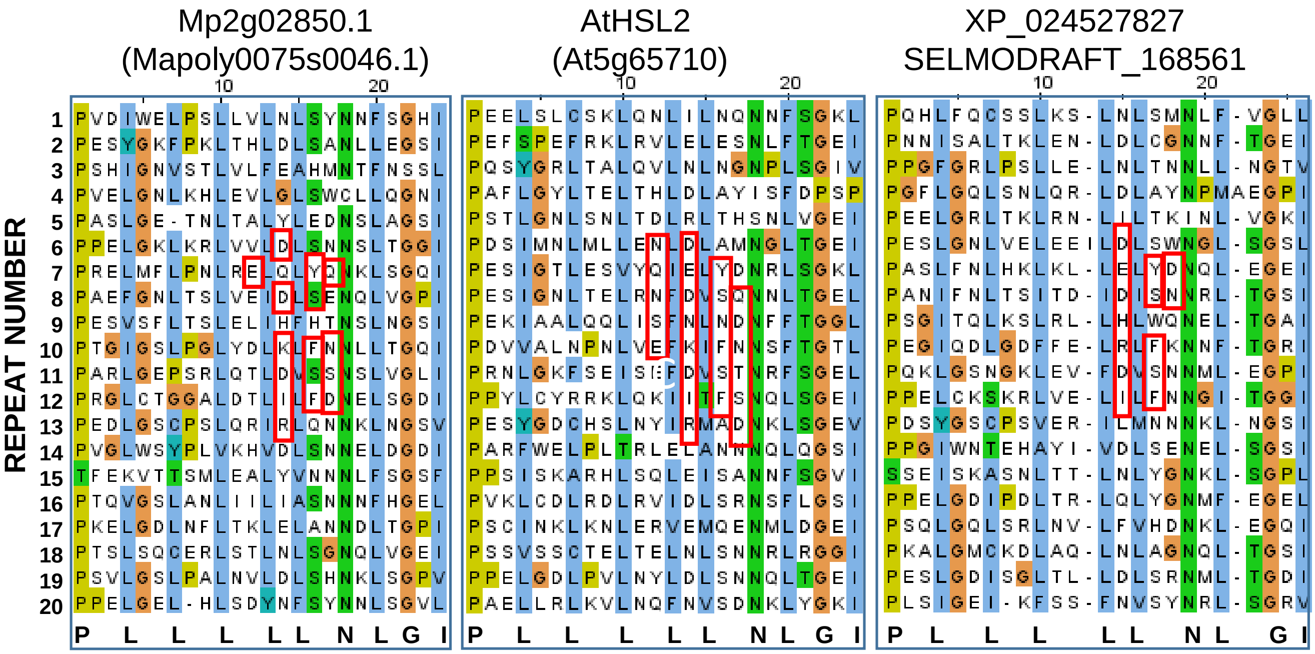


**Supplemental Figure 3.** Conservation of amino acids residues involved in ligand binding. Support for Figure 4 and 6.

Alignment of the 24 amino acids long leucine-rich repeat units of AtHSL2 and its putative orthologues of *M. polymorpha* and *S. moellendorffii.* Residues involved in ligand binding according to the crystal structure of the receptor with the IDA peptide (Santiago et al., 2016) are marked with red. rectangles The position of amino acids making up the LRR scaffold are show at the bottom of the alignments. Residues are color-coded based on their chemical characteristics.

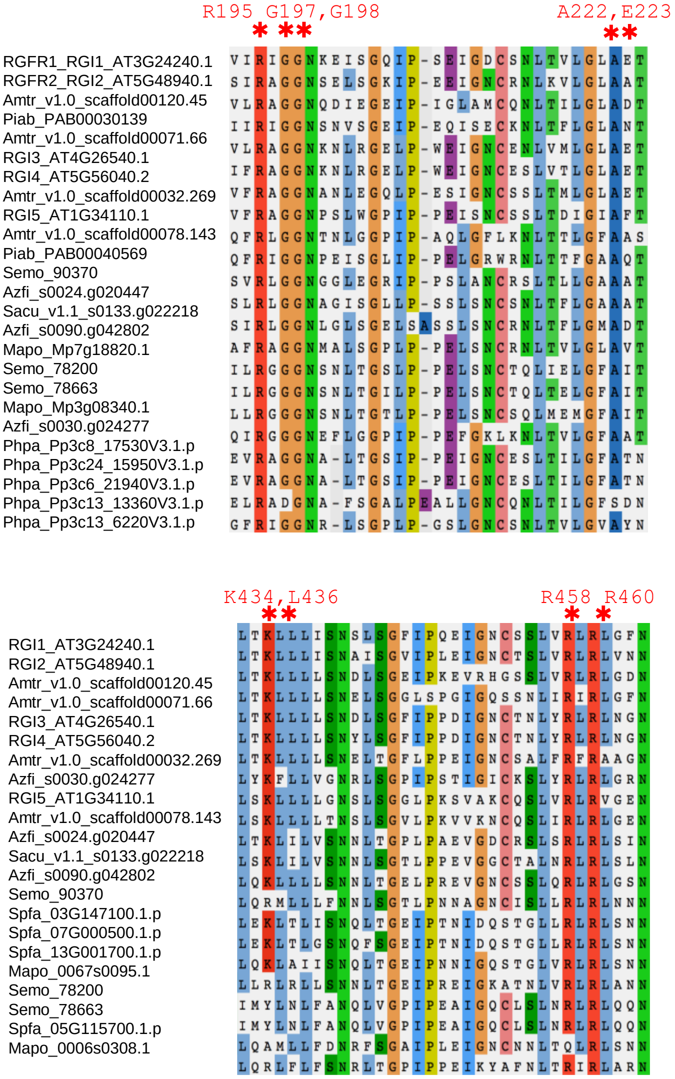


**Supplemental Figure 4.** Conservation of ligand-binding residues in RGI receptors. Support for Figure 5.

Alignments of RGF-binding regions of the LRRs of RGI homologues from diverse species. Residues shown to bind according to crystal structure (Song et al., 2016) are shown in red above the alignments.

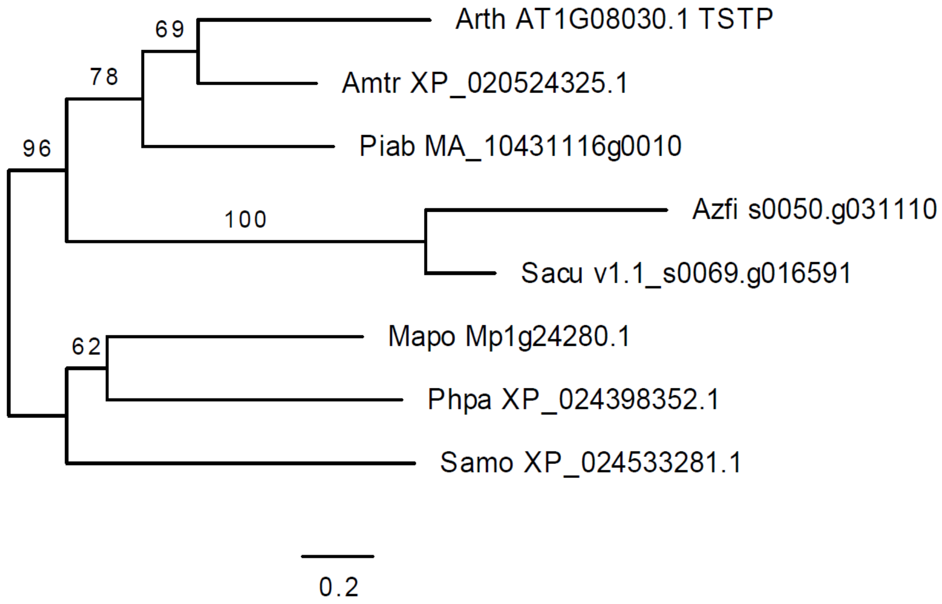


**Supplemental Figure 5.** Phylogenetic analysis of TYROSYLPROTEIN SULFOTRANSFERASE (TPST). Support for Figure 5.
Phylogenetic tree based on the ectodomain amino acid sequence of Arabidopsis TPST and orthologues in *A. trichopoda,* the gymnosperm *P. abies*, the ferns *A. filiculoides* and *S. cucullata,* the lycophyte *S. moellendorffii, the moss P. patens,* and the bryophyte *M. polymorpha* constructed by the maximum likelihood method. Bootstrap values are indicated at the nodes as percentages. The tree is drawn to scale, with branch lengths measured as the number of substitutions per site.
